# Identification of a broad and potent V3 glycan site bNAb targeting an N332_gp120_ glycan-independent epitope

**DOI:** 10.1101/2025.09.05.674437

**Authors:** Lutz Gieselmann, Andrew T. DeLaitsch, Malena Rohde, Caelan Radford, Johanna Worczinski, Anna Momot, Elvin Ahmadov, Judith A. Burger, Colin Havenar-Daughton, Sharvari Deshpande, Federico Giovannoni, Davide Corti, Christoph Kreer, Meryem Seda Ercanoglu, Philipp Schommers, Ivelin S. Georgiev, Anthony P. West, Jacqueline Knüfer, Ricarda Stumpf, Arne Kroidl, Christof Geldmacher, Lucas Maganga, Wiston William, Nyanda E. Ntinginya, Michael Hoelscher, Zhengrong Yang, Qing Wei, Matthew Renfrow, Todd J. Green, Jan Novak, Marit J. van Gils, Harry B. Gristick, Henning Gruell, Jesse D. Bloom, Michael S. Seaman, Pamela J. Bjorkman, Florian Klein

## Abstract

Broadly neutralizing antibodies (bNAbs) against HIV-1 can suppress viremia *in vivo* and inform vaccine development. Here, we characterized 007, a V3 glycan site bNAb exhibiting high levels of antiviral activity against multiclade pseudovirus panels^1–3^ (GeoMean IC_50_ = 0.012 µg/mL, breadth = 69%, 217 virus strains) by targeting a N332_gp120_ glycan-independent V3 epitope, a site of Env vulnerability to which only weakly neutralizing antibodies had previously been identified. Functional analyses demonstrated distinct binding and neutralization profiles compared to classical V3 glycan site bNAbs. A 007 Fab-Env cryo-EM structure revealed contacts with the V3 ^324^GD/NIR^327^ motif and interactions with N156_gp120_ and N301_gp120_ glycans. In contrast to classical V3 bNAbs, 007 binding to Env does not depend on the N332_gp120_ glycan, rendering it resistant to common escape mutations. Structures of 007 IgG-Env trimer complexes showed two Env trimers crosslinked by three bivalent IgGs, and bivalent 007 IgG was up to ∼300-fold more potent than monovalent 007 IgG heterodimer, suggesting a role for avidity in potent neutralization. Finally, in HIV-1_ADA_-infected humanized mice, 007 caused transient decline of viremia and overcame classical V3 escape mutations, highlighting 007’s potential for HIV-1 prevention, therapy, functional cure, and vaccine design.

## Introduction

Broadly neutralizing antibodies (bNAbs) targeting the HIV-1 envelope protein (Env) inform vaccine design, hold potential for therapy and prevention, and advance efforts toward achieving a functional cure^4–9^. However, clinical trials have underscored the stringent requirements for enhanced antiviral activity that will be critical to counteract virus Env diversity and emergence of escape. Combined application of bNAbs with complementary neutralization coverage offers an opportunity to overcome these challenges^10–14^. Thus, the discovery of new bNAbs demonstrating distinct binding modes, neutralizing profiles, and viral escape pathways remains essential to facilitate successful clinical application of bNAbs.

On the HIV-1 Env trimer, bNAbs recognize highly conserved epitopes essential for viral entry^15–20^. One such epitope is a V3 glycan site located at the base of the V3 loop. This epitope includes the N332_gp120_ glycan, the ^324^GD/NIR^327^ motif, and N-linked glycans in the vicinity (N133_gp120_, N137_gp120_, N156_gp120_, and N301_gp120_)^21–23^. However, some V3 glycan site bNAbs are also known to exhibit promiscuity in their glycan recognition and/or accommodation, allowing tolerance to shifts in N-glycan composition and configuration. For example, while bNAbs such as 10-1074, BG18, PGT124, and DH270 are highly dependent on the presence of the N332_gp120_ glycan, others like PGT121, PGT128 and PGT130 can compensate for the loss of the N332_gp120_ glycan by targeting alternative glycans within the high-mannose patch^24–29^. Recently, a neutralizing antibody (nAb), EPTC112, that lacks contacts with the N332_gp120_ glycan and instead targets a previously undescribed N332_gp120_ glycan-independent V3 epitope extending to glycans of the V1 loop was reported^30^. However, EPTC112 displayed low levels of breadth (25%, cut-off ≤10µg/mL, 129 virus strains)^31^ and potency (GeoMean IC_50_ against all strains = 3.7 µg/mL)^31^, limiting its applicability for vaccine design, prevention and immunotherapy.

Anti-HIV-1 bNAbs neutralize the virus through multiple mechanisms including blocking of receptor binding, hindering membrane fusion, and accelerating decay of Env trimers^15^. As with antibodies to other antigens^32^, IgG bivalency could enhance the breadth and potency of HIV-1 bNAbs by permitting the binding of adjacent Envs on the surface of the virus (inter-spike crosslinking) or by simultaneously engaging with multiple protomers of the same Env (intra-spike crosslinking)^33,34^. However, HIV-1 bNAb IgGs are usually not known to utilize bivalent binding, a likely consequence of the relatively few Envs coating the surface of the virus and the positioning of conserved bNAb epitopes preventing inter- and intra-Env crosslinking, respectively^33^. Thus, the discovery of naturally occurring bNAbs that utilize avidity presents an opportunity for optimizing prevention and immunotherapy, as well as for informing vaccine design.

Here, we report on the identification and detailed characterization of the new anti-HIV-1 bNAb 007 targeting a N332_gp120_ glycan-independent V3 epitope of the Env trimer. Cryo-EM analyses revealed a distinct binding mode compared to canonical V3 glycan site bNAbs that depends on the N301_gp120_ and N156_gp120_ glycans. Bivalent 007 IgG was more potent than its monovalent forms, and a structure of 007 IgG in complex with SOSIP trimers revealed a stable dimer of Env trimers linked by IgGs. Moreover, 007 displayed high levels of antiviral activity and exhibited a different neutralization profile compared to V3 reference bNAbs against extended multiclade panels. In HIV-1_ADA-_infected humanized mice, 007 achieved transient decline of viremia and overcame V3 escape mutations. In conclusion, our findings suggest the N332_gp120_-independent V3 epitope as a viable target for vaccine design and expand the armamentarium of bNAbs available for clinical use.

## Results

### Identification of bNAb 007 with distinct binding and neutralizing profile

Donor EN01 was identified as the top elite neutralizer from Mbeya region in southern Tanzania which was part of a multinational cohort of 2,354 people living with HIV (PLWH)^35^. Purified serum immunoglobulin G (IgG) from donor EN01 displayed the highest levels of breadth (100%) and potency (mean neutralization of 99% at 300 µg/ml) against the 12-strain global HIV-1 pseudovirus panel^3,35^ (Fig. 1a, Extended Data Fig. 1a). Neutralization fingerprint analyses to determine the potential epitope specificity of the serum IgG response were inconclusive due to ambiguous delineation scores^35,36^ (Extended Data Fig. 1b). To identify nAbs mediating the broad and potent serum response, we isolated single HIV-1 Env-reactive B cells using a GFP-labeled BG505_SOSIP.664_ bait protein^37^. HIV-1 Env-reactive B cells were isolated at a frequency of 0.46% and a total of 189 heavy chain and 100 light chain sequences were amplified using optimized PCR primers and protocols^38,39^ (Extended Data Fig. 2a). Heavy chain sequence analyses revealed a high degree of clonality among the isolated BCR sequences, identifying 23 B cell clones with two or more members. Compared to a human reference memory IgG repertoire dataset^40^, amplified BG505_SOSIP.664_-reactive V_H_ sequences of donor EN01 displayed higher levels of somatic mutation (median V_H_ gene germline identity of 85.3% vs. 95.3% on nucleotide level), an enrichment of V_H_ gene segments 1-69 and 4-4, and comparable CDRH3 lengths (Extended Data Fig. 2a,b).

**Figure 1:**
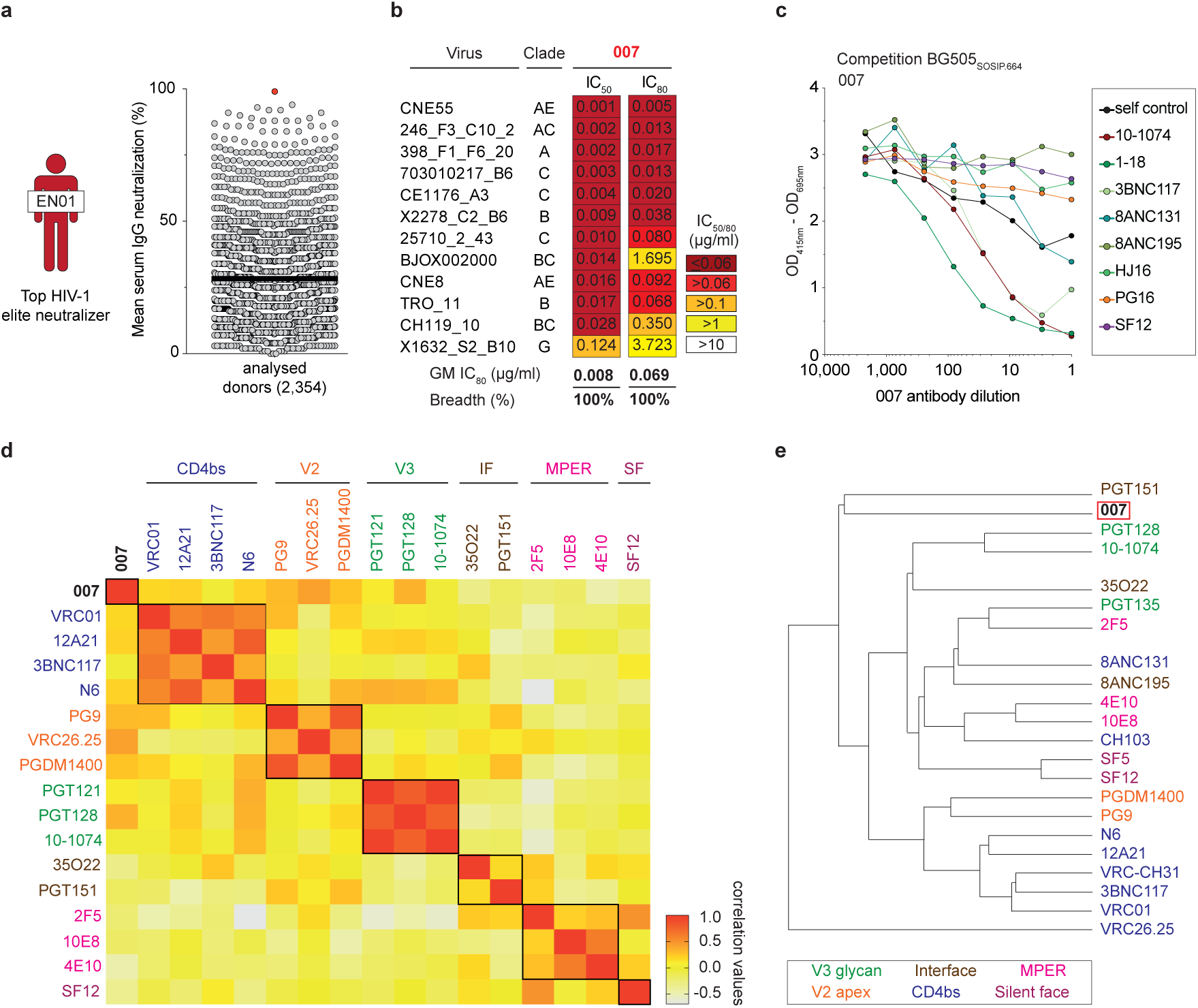
Identification of bNAb 007 with distinct binding and neutralizing activity. **a**, HIV-1 neutralizing serum activity of HIV-1 elite neutralizer EN01^112^ against the global^3^ and f61 fingerprinting^36^ pseudovirus panel. Serum IgG samples were tested in duplicates^35^. **b**, Neutralization activity of bNAb 007 against the HIV-1 global pseudovirus panel. Samples were tested in duplicates. **c**, Interference of 007 with selected reference bNAbs targeting known epitopes on the HIV-1 Env trimer, as determined by competition ELISAs. **d**, Comparison of the neutralization profile of 007 with reference bNAbs targeting known epitopes against the f61 fingerprinting panel and **e**, 119 multiclade pseudovirus panel^1^. Antibody 007 was tested in duplicates. Neutralization data of reference bNAbs (**b**, **d** and **e**) were retrieved from CATNAP^31^ database.

A total of 48 representative mAbs were produced and evaluated for neutralizing activity at a concentration of 2 µg/mL against a panel of six HIV-1 pseudoviruses representing different clades (Extended Data Fig. 2c). Among the screened mAbs, 007 identified from B cell clone 17 neutralized all strains of the screening panel with a mean neutralization of 90% at 2 µg/mL. Members of this clone derive from V_H_4-34*01/02 and V_K_1-12*01 gene segments with CDRH3 and CDRL3 lengths of 22 and 9 amino acids (Supplementary Table 1). V_H_ gene germline identities from this clonal family ranged from 75.0% to 80.5%, while V_K_ gene germline identities ranged from 76.2% to 85.0% at the nucleotide level (Supplementary Table 1). All members of B cell clone 17 exhibited high antiviral activity against the 12-strain global HIV-1 pseudovirus panel^3^, with 007 being the most potent antibody of the clone (Breadth=100%; GeoMean IC_50_/IC_80_=0.008/0.069 µg/mL) (Fig. 1b, Supplementary Table 2). To map the epitope of 007, we performed competition ELISAs using the BG505_SOSIP.664_ Env timer containing the T332N amino acid substitution. We detected interference with the CD4bs bNAbs 1-18^41^ and 3BNC117^42^ as well as with the V3 glycan site bNAb 10-1074^24^ (Fig. 1c). In addition, we determined the neutralizing activity of 007 against BG505_T332N_ pseudovirus mutants that abrogate the activity of V3 glycan site^24^, CD4bs^43–46^, V2 apex^47^, MPER, and silent face^48^ reference bNAbs. None of the tested signature escape mutations reduced the neutralizing activity of 007 (Extended Data Fig. 2d). To delineate the neutralization profile of 007, we evaluated the antiviral activity against the f61 fingerprint^36^ and 119 multiclade pseudovirus panel^1^ (Extended Data Fig. 2e, Supplementary Table 3). Against these panels the neutralizing activity of 007 did not show a consistent correlation or alignment with reference bNAbs from known epitope classes (Fig. 1d,e). Next, we evaluated the potential reactivity of 007 to self-antigens using a HEp-2 staining assay^49^. Unlike the anti-HIV-1 bNAb 2F5, which exhibits known autoreactivity^50^, 007 showed no detectable autoreactivity against HEp-2 cells (Extended Data Fig. 3a). However, 007 demonstrated a pharmacokinetic profile less favorable to that of bNAbs 10-1074 and 3BNC117, which serve as reference for IgG1 half-lives in humans^51^ (Extended Data Fig. 3b). These findings indicate that 007 achieves high antiviral activity without evidence of autoreactivity and targets an epitope distinct from previously characterized bNAbs.

### 007 targets a N332_gp120_ glycan-independent V3 epitope on Env

To elucidate the binding mode of 007, we performed single-particle cryo-EM analysis of the antibody Fab in complex with a soluble BG505 Env trimer, which included SOSIP stabilizing mutations^52^ and an engineered disulfide (I201C_gp120_-A433C_gp120_) designed to prevent trimer opening (BG505-DS)^53,54^. Fab was added in a greater than 3-fold molar excess to SOSIP trimer, and Fab-Env complexes were then isolated by size-exclusion chromatography (SEC). Structural analysis revealed four trimer classes with either 0, 1, 2, or 3 bound Fabs, with the most populated class being a 3.0 Å single Fab-bound trimer (Fig. 2a, Extended Data Fig. 4, Supplementary Table 4). Comparison of the 007 binding pose to poses of V3 glycan site bNAbs showed a distinct mode of binding, most closely resembling the V3 glycan site bNAb PGT128 (Fig. 2b).

**Figure 2:**
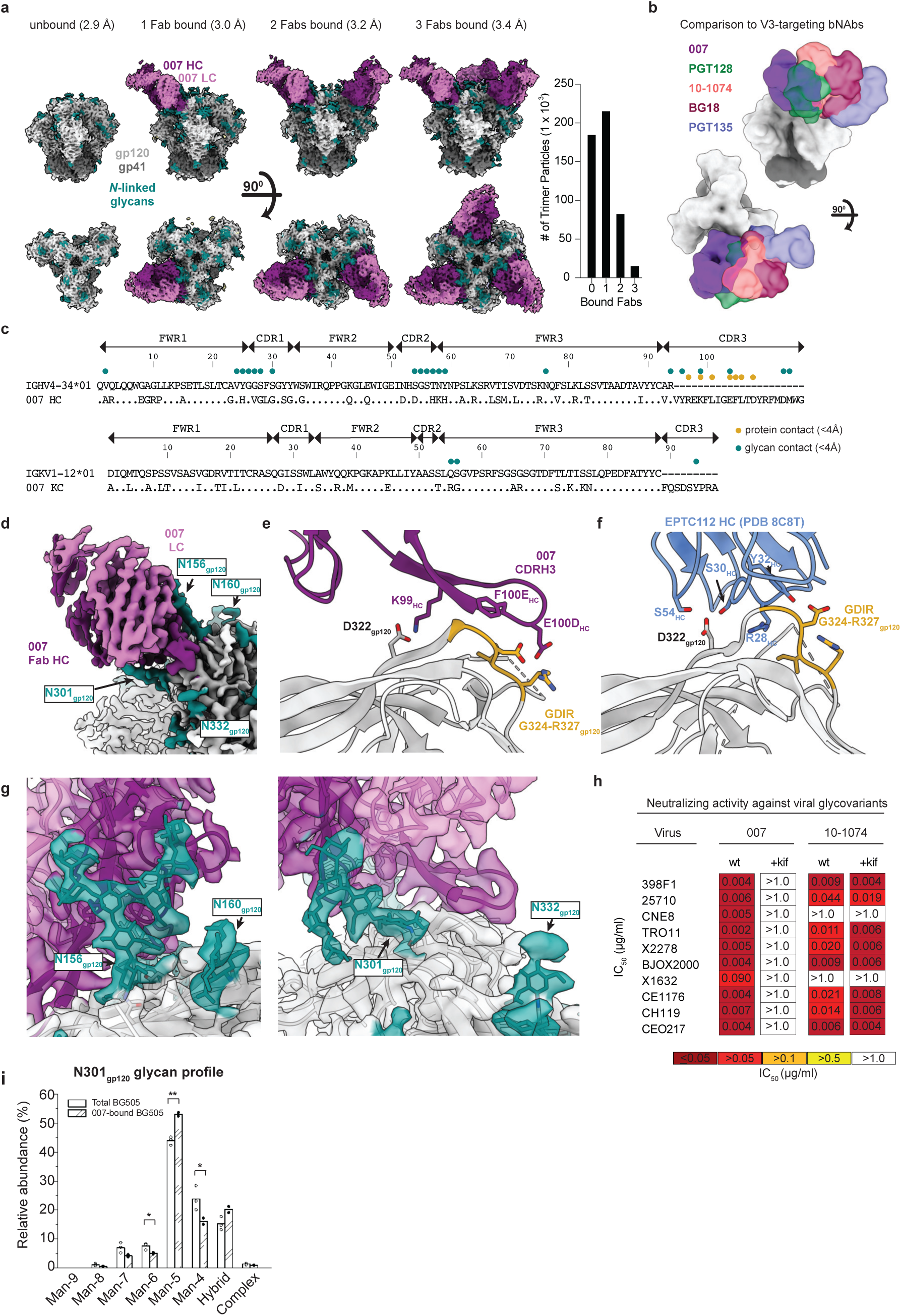
007 recognizes the N332_gp120_ glycan-independent V3 epitope. **a**, (left) Overviews of the four structural classes identified of SOSIP trimers with 0-, 1-, 2-, or 3-bound 007 Fabs per trimer. (right) The number of particles used in each of the final reconstructions. **b**, Overlay of 007 with V3-targeting bNAbs (PDB codes 5C7K, 5T3Z, 6CH7, 4JM2). **c**, Alignment of 007 VH and VL to their predicted germline V gene segments. 007 residues within 4Å of protein or glycan components on Env are indicated by colored circles. **d**, Structure overview, highlighting proximal glycans. **e**, Protein contacts between 007 and Env. **f**, Protein contacts between EPTC112 and Env (PDB code 8C8T). **g,** EM density highlighting the N156_gp120_ (left) and N301_gp120_ (right) glycans. **h,** Neutralizing activity (IC_50_ and breadth of 007 and 10-1074) against HIV-1 pseudoviruses produced in the presence of kifunensine. **i,** Comparison of glycoform abundance between total BG505 SOSIP and 007-bound SOSIP at N301_gp120_ determined by LC-MS/MS, with asterisks denoting level of significance (* denotes 0.01 < P ≤ 0.05, ** denotes P ≤ 0.01).

The structure revealed 007 is framed by N-linked glycans attached to N156_gp120_ and N301_gp120_, but does not contact the glycan at N332_gp120_, which is important for binding of many V3 glycan bNAbs^26,55,56^ (Fig. 2c,d). Antibody contacts with protein portions of Env are mediated exclusively by a 22-residue CDRH3 which extends outward to contact the conserved ^324^GD/NIR^327^ motif on Env (Fig. 2c,e). Within this region, F100E_HC_ contacts G324_gp120_, and E100D_HC_ at the tip of the CDRH3 is in close proximity to R327_gp120_ (Fig. 2e). Other V3 bNAbs possess negatively-charged glutamate residues within their CDRH3s that may form electrostatic contacts with the positively-charged R327_gp120_ (E100I_HC_ in 10-1074^55^, PGT122^56^, and PGT124^57^ and E100G_HC_ in BG18^26^). In addition, K99_HC_ of 007 forms an electrostatic interaction with D322_gp120_ (D321(1)_gp120_ in some numbering nomenclatures) within V3 (Fig. 2e, Extended Data Fig. 5a), a residue that is a negatively-charged Asp or Glu in >70% of sequences (www.hiv.lanl.gov and^58^). The CDRH3 is also positioned near the gp120 V1 loop, which is located adjacent to V3, thereby allowing 007 L100A_HC_ and L100F_HC_ to contact R151_gp120_ within V1 (Extended Data Fig. 5b). Interestingly, in gp120 protomers bound by 007, but not in unbound gp120s, a portion of the V1 loop is disordered (Extended Data Fig. 5c) suggesting that 007 binding destabilizes the V1 loop.

The V1/V3 epitope of 007 resembles that of the recently described bNAb EPTC112^30^. Both antibodies depend on the N156_gp120_ and N301_gp120_ glycans, but not the N332_gp120_ glycan, and both contact the ^324^GD/NIR^327^ motif on Env. However, 007 exhibits greater breadth and is more potent (GeoMean IC_50_/IC_80_ = 0.01/0.03 vs. 0.32/0.29 µg/mL, breadth = 66/53% vs 27/14%, 110 or 105 virus strains) than EPTC112 (Supplementary Table 3). Structurally, EPTC112 contacts extend only to D325_gp120_ of the ^324^GD/NIR^327^ motif, whereas 007 interacts with the entirety of the motif (Fig. 2e,f). More extensive contact with the ^324^GD/NIR^327^ motif and better accommodation of the N156_gp120_ and N301_gp120_ glycans could contribute to the enhanced neutralization properties of 007 compared with EPTC112.

N-glycans comprise much of the 007 epitope (Fig. 2c,d). The FWRH1/CDRH1 of 007 contains a glycine-rich stretch of amino acids (GVHGVGLGGSGWG; G23_HC_ – G35_HC_), which wrap around the core pentasaccharide of the N156_gp120_ glycan (Extended Data Fig. 5d. Five of these glycine residues arose from somatic hypermutation (Fig. 2c), but only one is an improbable mutation, as defined by the ARMADiLLO web server^59^. A similar mechanism of glycan accommodation is utilized by the VRC01-class CD4-binding site bNAbs, which either acquire deletions or glycine substitutions in CDRL1 to accommodate the N276_gp120_ glycan^60^. In addition, the 007 CDRH2 packs against the N301_gp120_ glycan (Extended Data Fig. 5d). Although extensive EM density was observed for both glycans, density corresponding to a core fucose, a component of complex-type glycans, was not observed, and more density was seen for the N156_gp120_ glycan than the N301_gp120_ glycan (Fig. 2g). A caveat of these observation is that single-particle cryo-EM analysis of glycans is limited due to their compositional and conformational heterogeneity^61^.

To evaluate the role of N-glycan processing in 007 neutralization, we expanded our neutralization analysis to include pseudoviruses produced in the presence of kifunensine, an inhibitor that prevents processing of high-mannose N-glycans to complex-type carbohydrates^62^. In contrast to other V3-targeting bNAbs, but in common with EPTC112^30^, 007 did not neutralize kifunensine-treated viruses (Fig. 2h, Supplementary Table 5), implying that the Man-9 (i.e., Man9(GlcNAc)2) high-mannose glycan trimming by mannosidase I is required for potent neutralization. We also performed site-specific N-glycan analysis using quantitative mass spectrometry profiling to determine the specific glycoforms enriched in 007-bound BG505 SOSIPs compared to total BG505 SOSIPs. The potential N-linked glycosylation site (PNGS) at N301_gp120_ in 007-bound BG505 showed an increase in Man-5 high-mannose glycans compared to the same site in the total BG505 sample (Fig. 2i), consistent with EM density and neutralization data (Fig. 2g,h, Supplementary Table 5). Unambiguous glycoform identification at position N156_gp120_ was not possible, however, as both N156_gp120_ and N160_gp120_ PNGSs resided on the same glycopeptide (Extended Data Fig. 6).

### 007 bivalency is required for potent neutralization

The observation of sub-stoichiometric trimer binding by a bNAb Fab could result from a weak monovalent binding interaction, which is unexpected for a potent bNAb such as 007. A difference between structural and neutralization experiments, however, is the use of monovalent antibody Fabs for structural studies, as opposed to bivalent IgGs for neutralization assays. Bivalent IgG binding can compensate for a weak Fab antigen-binding affinity; however, HIV-1 bNAb IgGs generally do not utilize avidity due to the relatively few Env trimers coating the virus and the positioning of conserved bNAb epitopes on Env, thus limiting inter- and intra-spike crosslinking, respectively^33^. We have used molar neutralization ratios (MNRs) [IC_50_ Fab (nM)/IC_50_ IgG (nM)] to evaluate potential contributions of avidity in neutralization, noting that viruses with densely-packed spikes can exhibit MNRs over 1000, whereas MNRs for anti-HIV-1 bNAbs tend to be low^33^. Notable exceptions include V3 bNAbs PGT128^63^ and EPTC112^30^, which target similar Env epitopes as 007 (Fig. 2b,f) and were reported to be ∼30 to 2000-fold more potent when formatted as an IgG than as a Fab against viruses tested. Although the mechanism underlying the enhanced IgG potencies was not reported, it was speculated to result from inter-spike crosslinking between adjacent Env trimers on the virion surface.

To investigate the possibility that 007 IgG utilizes avidity during neutralization of pseudovirions, we repeated *in vitro* neutralization assays to compare Fab and IgG forms of 007, finding Fab/bivalent IgG MNRs >1000 for most viruses tested (Fig. 3a,b, Extended Data Fig. 7,a, Supplementary Table 6). As this comparison does not control for steric effects favoring IgG neutralization due to increased mass compared with a Fab (∼150kDa versus ∼50kDa), we also created bispecific 007 IgGs^64^ in which one Fab arm was replaced with the anti-CD3 antibody OKT3^65^, which does not recognize HIV-1 Env. We found that bispecific 007/OKT3 IgG was more potent than 007 Fab, with an average Fab/bispecific IgG MNR of ∼20 against all viruses tested (Fig. 3a,b, Extended Data Fig. 7,a, Supplementary Table 6), consistent with neutralization enhancement due to steric effects. However, bivalent 007 IgG was more potent than bispecific 007/OKT3 IgG, consistent with avidity effects that enhanced neutralization: bispecific IgG/bivalent IgG MNRs ranged from <10 against CEO217 and CNE8 viruses to >200 against X2278 and CH119 (Fig. 3a,b, Extended Data Fig. 7,a, Supplementary Table 6). Such variation is expected since enhancements due to avidity depend on multiple factors including the association and dissociation rates of a Fab for an Env antigen^66^, which are expected to vary for different viral strains.

**Figure 3:**
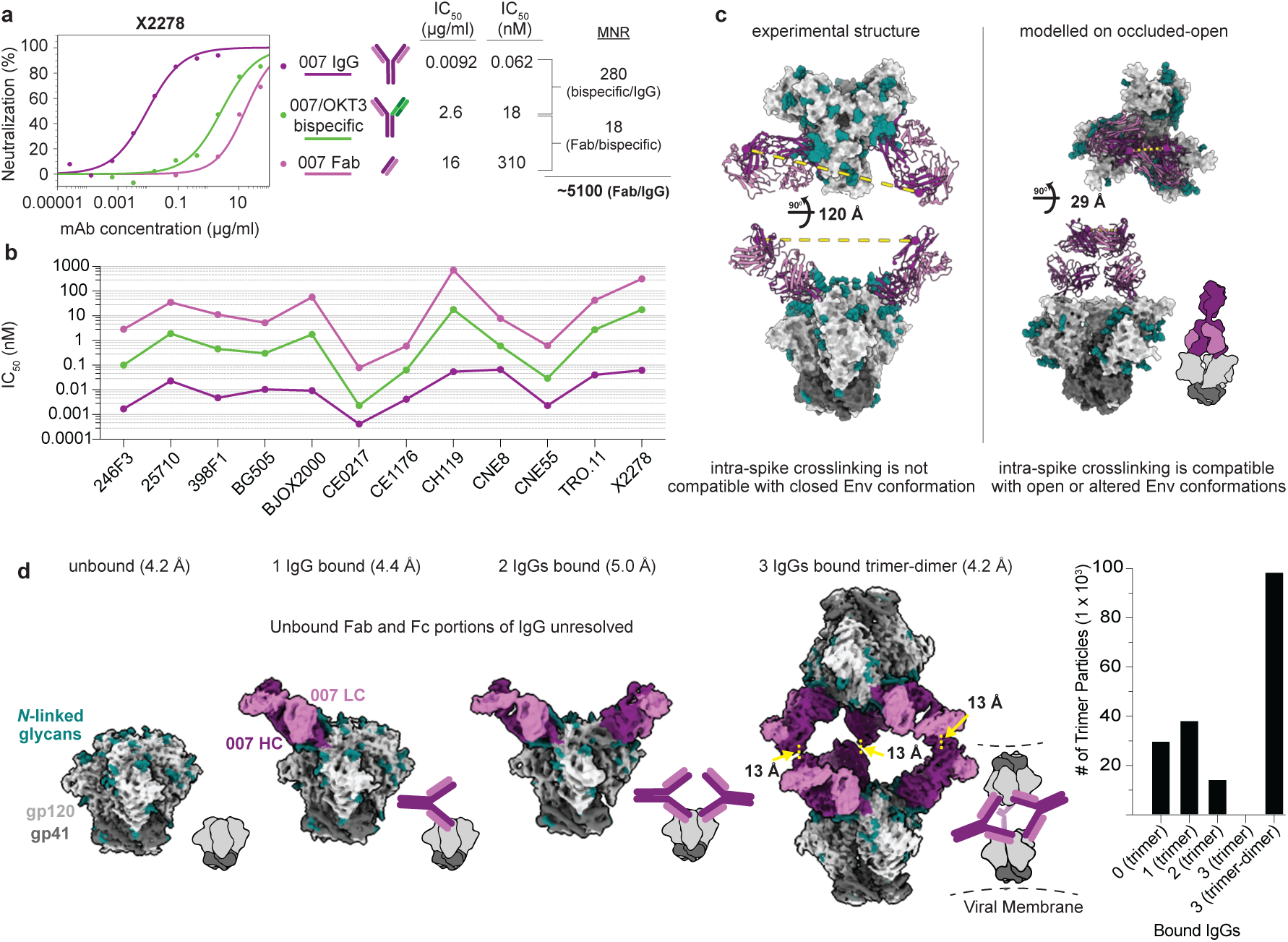
Potent neutralizing activity requires 007 bivalency. 007 exhibits avidity. **a**, Neutralization curves against strain X2278 (left) used to calculated molar neutralization ratios (MNRs) comparing the IC_50_ values for the IgG, bispecific, and Fab forms of 007 (right). **b**, Molar IC_50_ values for IgG, bispecific, and Fab forms of 007 against a panel of HIV-1 strains. Complete neutralization curves and IC_50_ values are in Extended Data Fig. 7a and Supplementary Table 6. **c**, Distance measurements between the C-termini of the Fab heavy chains on the experimentally determined 2 Fab-bound trimer structure (left) compared to a structure in which 007 Fab was modelled onto an occluded-open trimer conformation (PDB code 5VN8). A schematic showing a 007 IgG-bound occluded-open trimer is shown for clarity. **d**, (left) Four structural classes identified after complexing 007 IgG with BG505 SOSIP. Schematics are included for clarity. Dashed line (yellow) indicates the distance measurements between C-termini of Fab regions in 007 IgG-bound trimer-dimer structure. (right) The number of particles used in each of the final reconstructions. The number of particles in the trimer-dimer class were multiplied by two, as each particle contained two SOSIP trimers.

A possible mechanism for IgG avidity effects is through intra-spike crosslinking (i.e., both Fab arms on a bivalent IgG engage with epitopes on a trimeric Env), which has been previously inferred from Fab-trimer structures for other viral pathogens: e.g., as described for anti-SARS-CoV-2 antibodies^67–69^, a measured distance < ∼65Å between the C-termini of adjacent Fab heavy chains raises the possibility for intra-spike crosslinking by an IgG, as this would permit the C-termini of two Fab heavy chains to come together to form the N-terminus of the IgG Fc region. In our structure of BG505-DS SOSIP with two copies of 007 Fab, the measured distance between the C-termini of adjacent Fab heavy chains, ∼120 Å, was ∼2-fold greater than this distance cutoff (Fig. 3c), suggesting that intra-spike crosslinking by 007 IgG interacting with the closed BG505 Env trimer in our structure would not be possible. However, one must also consider different structural states of the antigen that may be targeted. For example, double electron-electron resonance (DEER) spectroscopy demonstrated that unliganded SOSIP Env trimers can sample open states that can be recognized by neutralizing antibodies, such as the occluded-open Env trimer state in which the gp120 protomers are outwardly rotated but the V1/V2 loop is not displaced to the sides of the Env trimer^70^. Docking the 007 Fab-gp120 coordinates onto the gp120s of an occluded-open Env placed the C-termini of adjacent Fabs within 65 Å (Fig. 3c). Therefore, one possibility to account for the apparent involvement of avidity effects in 007 interactions with Env trimers is that the 007 IgG binds bivalently to an occluded-open or other altered Env conformation that was not observed in our 007 Fab-SOSIP complex structures.

In the context of SARS-CoV-2 spike trimers, structural studies of IgGs interacting with stabilized trimers revealed intra-spike crosslinking^71,72^. Thus, in an attempt to investigate bivalent IgG interactions with Env trimer structurally, we incubated BG505 SOSIP with 007 IgG and imaged by cryo-EM (Extended Data Fig. 8, Supplementary Table 4). Rather than observing intra-spike crosslinking, the most populated structural class of 007 IgG-BG505 complexes contained “trimer-dimers” in which two SOSIP Envs were crosslinked by Fabs from three IgG molecules (Fig. 3d). This assembly exhibited D3 symmetry, with the apexes of two Env trimers facing each other and separated by ∼70Å. Density for the Fc region of the IgG was not resolved, an expected consequence of flexibility at the IgG hinge region^73^. However, the distance between the C-termini of the closest Fabs on the opposing Env trimers was 13 Å (Fig. 3d), consistent with these densities originating from a single IgG. In addition to the trimer-dimer structural class, trimer classes with 0, 1, or 2 copies of bound 007 IgG were also observed (Fig. 3d). In these complex structures, the unbound Fab and Fc of each IgG were unresolved, leading to EM densities closely matching the densities of the Fab-SOSIP complexes (Fig. 3d, 2a).

The observed trimer-dimer structure would be compatible with Envs attached to separate virions (Fig. 3d), thus raising the possibility that the 007 IgG neutralizes at least in part by aggregating virions. However, the extent to which enhanced pseudovirus neutralization by 007 IgG can be attributed to viral aggregation, intra-spike crosslinking, or another mechanism such as inter-spike crosslinking^30,63^, heteroligation^74^, or by enhancing gp120 shedding^72^ warrants further investigation.

### 007 exhibits high levels of antiviral activity and a complementary neutralization profile to V3-specific bNAbs

To further characterize the neutralizing properties of bNAb 007 in comparison to other V3 loop-targeting bNAbs, we compared their activities against 108 common strains from the 119-strain multiclade pseudovirus panel consisting of difficult-to-neutralize (tier-2 and tier-3) viruses, major genetic HIV-1 subtypes, and circulating recombinant forms (CRFs)^1^ (Fig. 4a, Supplementary Table 3). Notably, bNAb 007 demonstrated superior breadth and/or potency compared to reference V3 glycan site bNAbs across this panel (GeoMean IC_50_/IC_80_=0.01/0.03 µg/mL; breadth=66%/54%). Moreover, 007 greatly exceeded the neutralizing activity of bNAb EPTC112^30^, a recently identified antibody of the same epitope class. In addition, 007 demonstrated high neutralization activity against a 100-strain clade C panel^2^ (GeoMean IC_50_/IC_80_=0.01/0.03 µg/mL; breadth=68%/49%) (Extended Data Fig. 9a, Supplementary Table 7). Analyses of the antiviral activity against the 119-strain multiclade panel revealed that 007 displays a distinct neutralization profile compared to V3 glycan site reference bNAbs. While reference V3 bNAbs only neutralized up to 50% of Clade AC (2 of 4 strains) and 67% of CRF AE viruses (10 of 15 strains), 007 achieved 100% and 80% breadth, respectively (Fig. 4b). Furthermore, unlike other V3 glycan site bNAbs, 007 maintained high neutralization coverage against viruses lacking the N332_gp120_ glycan (breadth= 71%) and those with amino acid substitutions in the ^324^GD/NIR^327^ motif (Breadth= 50%) (Fig. 4c). Consistent with structural analyses, the antiviral activity of 007 was dependent on the N156_gp120_ and N301_gp120_ glycans, with viruses lacking PNGSs at these positions being completely resistant to neutralization by 007 (Fig. 4c). Beyond virus neutralizing activity, 007 also demonstrated strong Fc effector functions against Env expressing cells: in both FcγRIIIa signaling and natural killer cell-mediated antibody dependent cytotoxicity (ADCC) assays, 007 exhibited greater potencies compared to reference bNAb 10-1074 (Extended Data Fig. 9b). Altogether, 007 exhibits a potent antiviral profile, via both neutralization and Fc effector function mechanisms.

**Figure 4:**
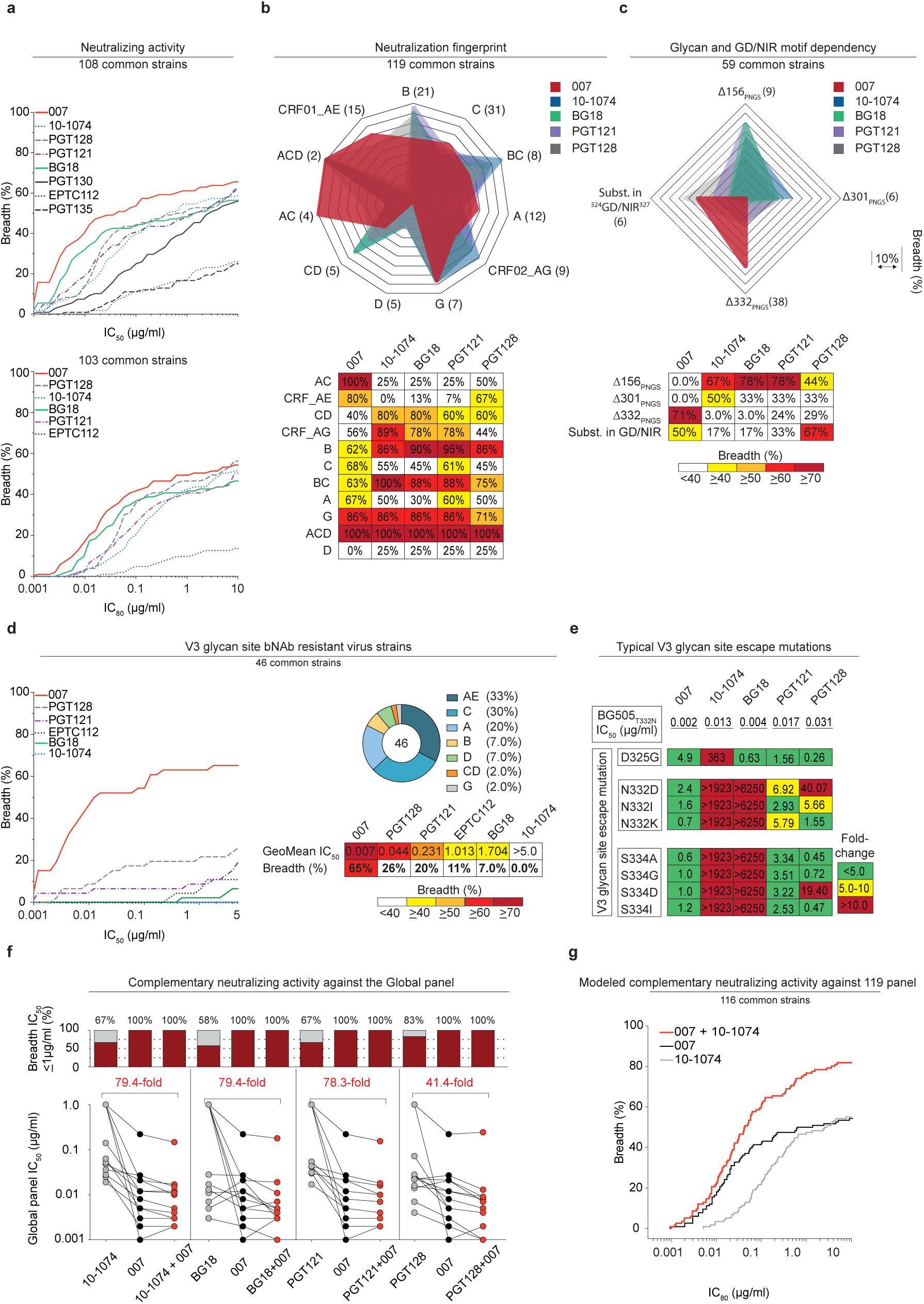
High antiviral activity against resistant virus strains driven by distinct neutralization profile and ^324^GD/NIR^327^ motif dependency. **a**, Neutralization breadth (%) and potency (IC_50_/IC_80_) of 007 against the 119 multiclade pseudovirus panel^1^. **b**, Illustration of the neutralization profile of 007 in comparison to V3 glycan site reference bNAbs against different virus clades of the 119 multiclade panel. **c**, Dependency of 007 and V3 glycan site reference bNAbs on potential N-linked glycosylation sites and the ^324^GD/NIR^327^ motif. **d**, Neutralizing activity (GeoMean IC_50_, breadth) of 007 compared with V3 glycan site reference bNAbs against a panel of 46 pseudovirus strains resistant to V3 glycan site bNAbs. Pie chart illustrates the clade distribution of resistant pseudovirus strains. **e**, Neutralizing activity against viral escape mutations within the V3 loop of the HIV-1 Env trimer. The top row displays bNAb IC_50_ values for the BG505_T332N_ pseudovirus, while panels illustrate changes in bNAb sensitivity (IC_50_ fold change) of pseudovirus mutants relative to BG505_T332N_. Antibodies were tested in duplicates. **f**, Neutralizing activity of 007 in combination with V3 glycan site bNAbs (mixed at a 1:1 ratio) against the global pseudovirus panel^3^. Single and combined mAbs were tested up to a concentration of 1µg/mL (total IgG amount). Red numbers indicate the fold change in IC_50_s (increase in potency) between the individual mAb and its combination with 007. **g,** Computational modeling of the predicted neutralizing activity of bNAb 007 in combination with 10-1074 against the 119 multiclade pseudovirus panel^1^ using the CombiNaber tool^14^ (http://www.hiv.lanl.gov/content/sequence/COMBINABER/combinaber.html). Breadth (%) was calculated using a cut-off of ≤10 μg/mL (**a**-**d**, **g**). Data are shown for identical virus strains across each panel with available reference neutralization data (**a**-**d**, **g**). Reference bNAb data were sourced from the CATNAP database^31^.

Given its distinct neutralization profile and glycan dependency, we further analyzed the antiviral activity of 007 against 46 pseudovirus strains selected from the 119-strain multiclade pseudovirus panel^1^. This subset represents diverse HIV-1 clades and exhibits full resistance to the classical V3 glycan site bNAb 10-1074. Notably, while V3 reference bNAbs neutralized only 0%– 26% of these virus strains, 007 achieved a breadth of 65% with high potency (GeoMean IC_50_ = 0.007 µg/mL) (Fig. 4d). To further investigate how known Env escape mutations influence 007’s neutralizing activity, we evaluated the sensitivity of HIV-1_BG505_ pseudovirus variants^75^. Amino acid substitution at position D325_gp120_ and removal of the PNGS at position N332_gp120_ had variable effects on the activity of V3 reference bNAbs (Fig. 4e). Whereas the neutralizing activities of 10-1074 and BG18 were abrogated by all tested mutations at N332_gp120_ and S334_gp120_, only N332D_gp120_ and S334D_gp120_ greatly reduced PGT128 activity (Fig. 4e). Furthermore, among the V3 reference bNAbs tested, only 10-1074 was affected by D325G_gp120_, whereas PGT121 remained relatively unaffected by all assessed mutations (Fig. 4e). In contrast, 007 maintained robust antiviral activity against typical V3 glycan site escape variants^76^, showing minimal or no susceptibility to these mutations (Fig. 4e). Supporting these findings, the combination of 007 with other V3 glycan site bNAbs complemented their individual antiviral activity, leading to increased breadth and potency (>40-fold) of the combination against the global pseudovirus panel (Fig. 4f, Supplementary Table 8). In addition, *in silico* modeling of 007 and 10-1074 combination predicted complementary neutralizing activity, enhancing the breadth and potency against the 119 multiclade panel to 86.2% and a GeoMean IC_50_ of 0.011 µg/mL (Fig. 4g). The distinct neutralization profile of 007 establishes it as complementary to V3 glycan site bNAbs, offering broader coverage of viruses that evade neutralization by antibodies of this class.

### Deep mutational scanning reveals a distinct escape profile

Deciphering viral escape pathways is essential to inform clinical applicability of bNAbs. To more comprehensively investigate viral escape from bNAb 007, we employed a lentiviral pseudovirus-based deep mutational scanning (DMS) platform^77–79^ using two HIV-1 Envs from distinct clades: TRO.11 (Clade B)^80^ and BF520.W14M.C2 (Clade A)^81,82^ (Extended Data Fig. 5a). This platform encompasses nearly all functionally tolerated mutations at each individual Env residue, enabling a systematic evaluation of their effects on bNAb sensitivity and viral cell entry *in vitro*. DMS revealed a distinct viral escape profile for 007 compared to classical V3-targeting bNAbs. Specifically, mutations at the N332_gp120_ glycosylation site in both Env_TRO.11_ and Env_BF520_ conferred escape from classical V3-targeting bNAbs^24,25,83^. In contrast, 007 remained unaffected by these mutations (Fig. 5a, Extended Data Fig. 9c). Conversely, mutations that disrupted glycosylation at sites N156_gp120_, N188a_gp120_, and N301_gp120_ in Env_TRO.11_ reduced 007’s neutralizing activity but did not impair the activity of most other V3-targeting bNAbs. The only exception was PGT128, which was similarly affected by N301_gp120_ glycosylation loss (Fig. 5a). Due to functional intolerance of Env_BF520_ to mutations at N156_gp120_ and N301_gp120_, mutations at these sites could not be reliably assessed for this virus strain (Extended Data Fig. 9c).

**Figure 5:**
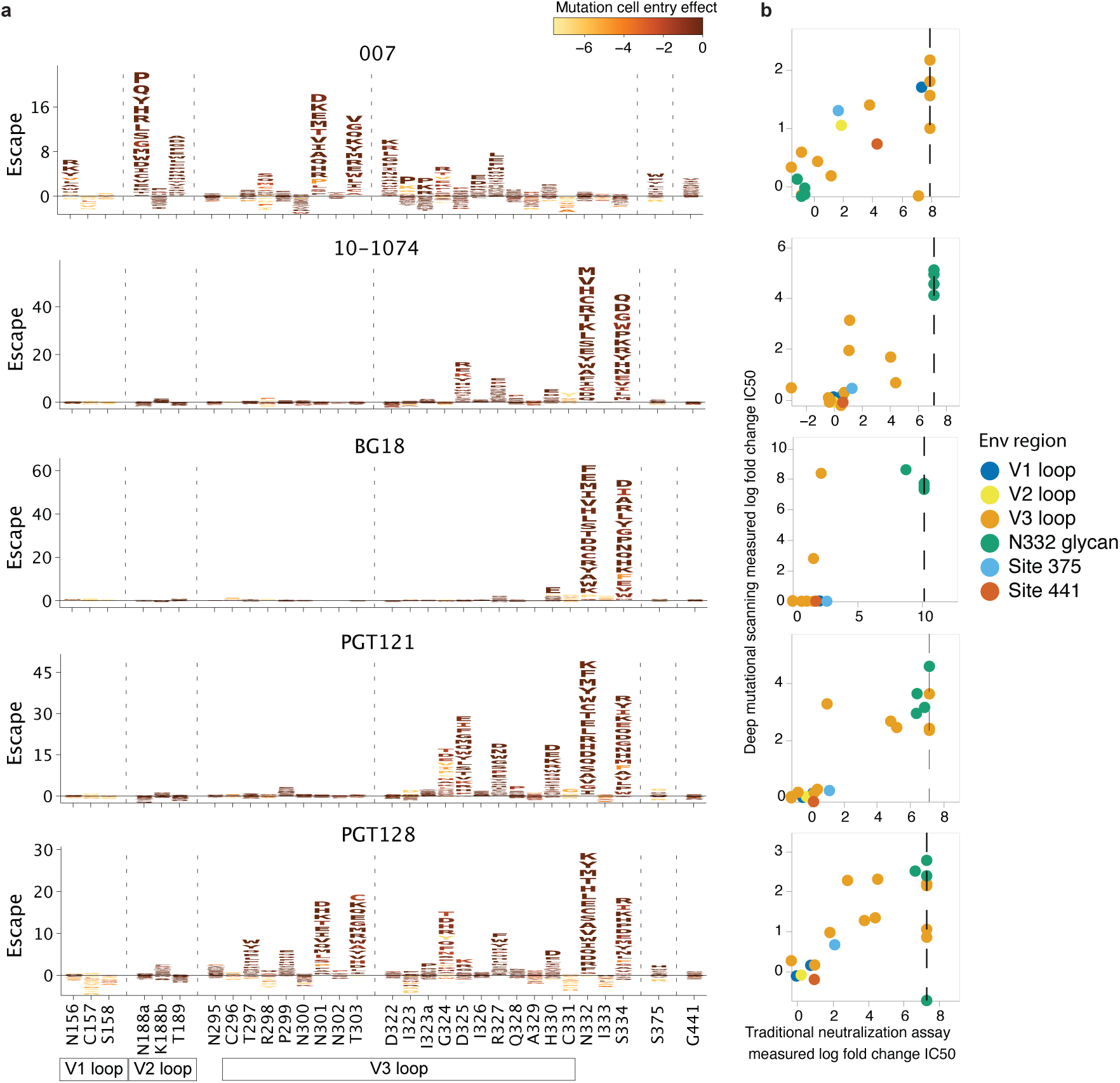
Deep mutational scanning analyses reveal distinct viral escape from bNAb 007 in HIV-1 Env_TRO.11_. **a**, Logo plots showing effects of mutations on neutralization escape in HIV Env_TRO.11_ for antibodies 007, 10-1074, BG18, PGT121, and PGT128. The height of each letter represents the effect of that amino-acid mutation on antibody neutralization, with positive heights (letters above the zero line) indicating mutations that cause escape, and negative heights (letters below the zero line) indicating mutations that increase neutralization. Letters are colored by the effect of that mutation on Env mediated cell entry function, with yellow corresponding to reduced cell entry and brown corresponding to neutral effects on cell entry. Only key sites are shown. See https://dms-vep.org/HIV_Envelope_TRO11_DMS_007/htmls/all_antibodies_and_cell_entry_overlaid.html for interactive versions of the escape maps that show all mutations. Escape maps against HIV-1 Env_BF520_ are shown in Extended Data Fig. 9c. **b**, Scatter plots of Env_TRO.11_ mutant fold-change IC_50_s measured by deep mutational scanning versus those measured in traditional neutralization assays. Each scatter plot shows log fold change IC_50_s for neutralization assays using the antibody labeling the logo plot in the same row. Each point represents the mean of two replicate neutralization curve measurements of one Env_TRO.11_ mutant. Env_TRO.11_ mutants are colored by Env region or site. Vertical dotted lines represent the limit of detection of the neutralization assays. See methods section “Deep mutational scanning data analysis” for details on how deep mutational scanning measured escape values are converted to deep mutational scanning measured IC_50_ values. The deep mutational scanning measured effects of mutations on escape from antibody 10-1074 shown in this Figure were previously published^77^.

In addition to escape due to substitutions in PNGSs, DMS uncovered distinct escape pathways from 007 at non-glycosylated residues in the V3 loop. While substitutions at Env_TRO.11_ residue R327_gp120_ facilitated escape from classical V3-targeting bNAbs, 007 was also affected by mutations at Env_TRO.11_ residues 322–323_gp120_ and Env_BF520_ residues 318–320_gp120_ (Fig. 5a, Extended Data Fig. 5a). Notably, substitutions introducing positive charges or eliminating negatively charged residues in this region (D322_TRO11_K/R or D321_BF520_K/H) mediated escape from 007, whereas introducing a negative charge in this region in BF520 (G322_BF520_D/E) enhanced neutralizing activity of 007, without impacting the neutralization capacity of other V3-targeting antibodies (Fig. 5a, Extended Data Fig. 9c), further highlighting the importance of the electrostatic interaction between K99_007_ and D322_gp120_ observed in the 007 structure with BG505.

To validate the viral escape pathways identified by DMS, we conducted TZM-bl neutralization assays^84^ incorporating the identified escape mutations in the HIV-1 Env_TRO.11_ background. The results revealed a high concordance between escape profiles identified by DMS and changes in neutralization potency measured in TZM-bl neutralization assays (fold changes in IC_50_s) (Fig. 5b Supplementary Table 9). These assays confirmed the pattern of viral escape observed in DMS analyses. Amino acid substitutions in the V1, V2 and V3 loop but not at the N332_gp120_ PNGS mediated escape from 007, setting this bNAb apart from classical V3-targeting antibodies.

### 007 suppresses viremia *in vivo* and dual-targeting of the V3 glycan site delays viral rebound

To investigate the *in vivo* activity of 007 and evaluate viral escape profiles, we investigated its effects in humanized mice infected with HIV-1_ADA_. Following a loading dose of 1 mg IgG, infected mice were treated with 0.5 mg of either 007 or 10-1074 IgG twice a week for up to 5 weeks (Fig. 6a, Extended Data Fig. 10). PBS-treated mice with matched stem cell donors served as controls, and log_10_ changes in viremia were normalized to this control group (Extended Data Fig. 10). Similar to 10-1074 IgG, administration of 007 IgG monotherapy resulted in a rapid decline in viral loads by 0.78 log_10_ copies/mL, followed by rebound of viremia within 14 days after treatment initiation (Fig. 6a). To characterize viral escape associated with rebound, we performed single genome sequencing (SGS) of plasma virus collected 5 weeks after treatment initiation and determined sensitivities to the corresponding bNAb (Fig. 6b, Supplementary Table 10). Viral escape from 10-1074 monotherapy was associated with mutations that abrogated the PNGS at N332_gp120_. In contrast, 13 env sequences from 4 mice receiving 007 monotherapy exhibited mutations at residues 139_gp120_, 146_gp120_, 303_gp120_,322_gp120_ and 341_gp120_ in the V1/V2 or V3 loop. Of these, only mutations at residues 303_gp120_ and 322_gp120_ mediated complete resistance to 007. Notably, all generated escape variants from 10-1074 monotherapy retained *in vitro* sensitivity to 007, and vice versa (Fig. 6B, Supplementary Table 10). We conclude that bNAb 007 reduces viremia *in vivo* and exerts a distinct and less restricted selection pressure on HIV-1 compared to 10-1074.

**Figure 6:**
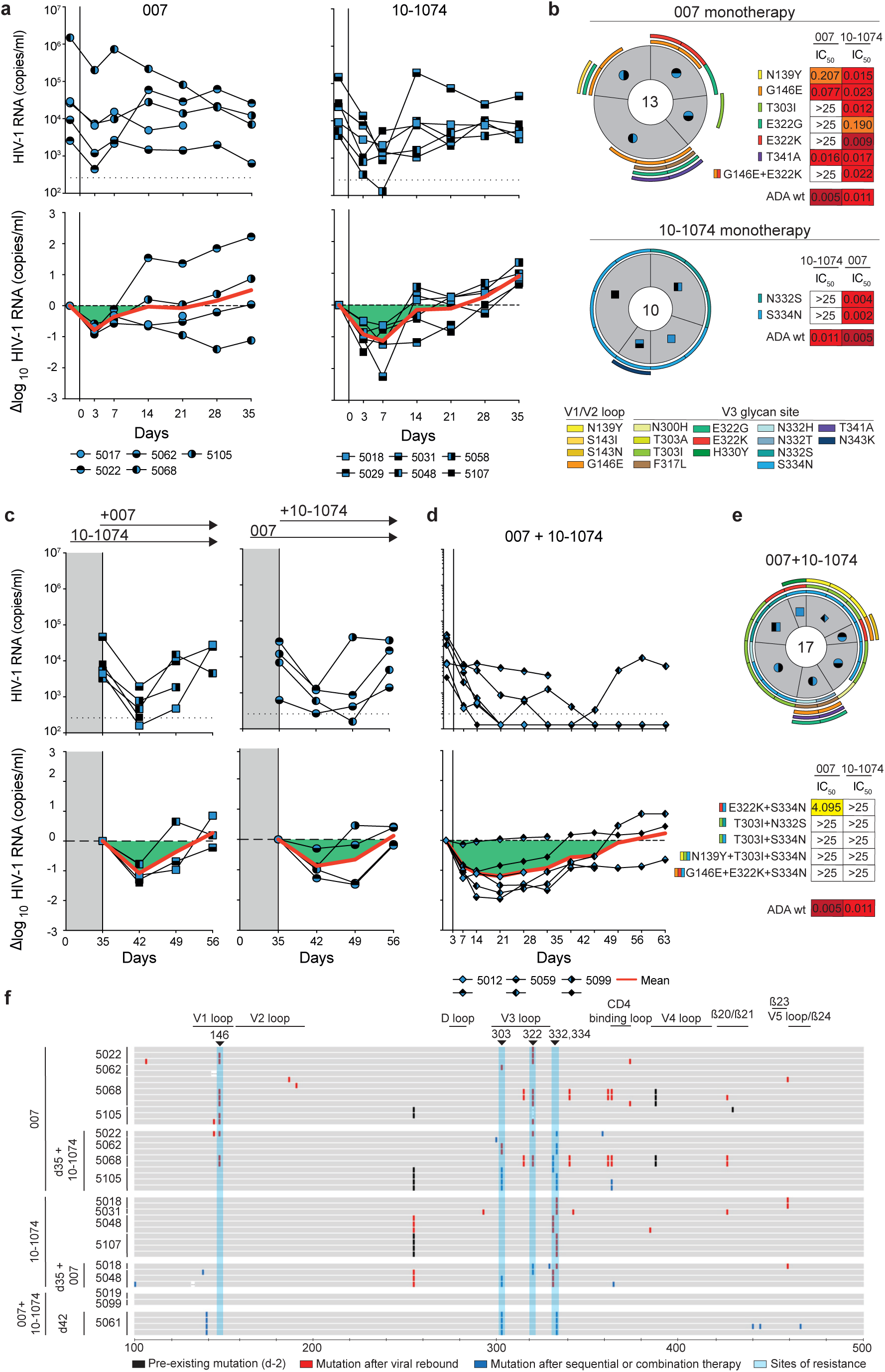
007 monotherapy and dual-targeting V3 glycan site combination therapy in HIV_ADA_-infected humanized mice. **a**, Investigation of the antiviral activity of 10-1074 and 007 monotherapy in HIV-1_ADA_-infected humanized mice. Graphs display the absolute HIV-1 RNA plasma copies/mL (top) and relative log_10_ changes from baseline viral loads (bottom) after initiation of bNAb therapy. Dashed lines (top graphs) indicate the lower limit of quantitation of the qPCR assay (LLQ) (260 copies/mL). Red lines display the average log_10_ changes compared to baseline viral loads (day −2). **b**, Analyses of single HIV-1 plasma env sequences from HIV-1_ADA_-infected humanized mice obtained after viral rebound on day 35 for 007 and 10-1074 monotherapy groups. Total number of analyzed sequences is indicated in the center of each pie chart. Mice are labeled according to icon legends in A. Colored bars on the outside of the pie charts indicate mutations in V1/V2 loop and V3 loop. Sensitivity (IC_50_s) of pseudoviruses generated from SGS-derived sequences against 007 and 10-1074. **c**, Sequential treatment with 007 or 10-1074 in HIV-1_ADA_-infected humanized mice following viral rebound during 007 or 10-1074 monotherapy (from panel **a**). This approach included maintaining 007 or 10-1074 monotherapy while integrating 007 or 10-1074 in the treatment regimen. Dashed lines (top graphs) indicate the lower limit of quantitation of the qPCR assay (LLQ) (260 copies/mL). Red lines display the average log_10_ changes compared to baseline viral loads (day 35). **d**, Antiviral activity of 007 and 10-1074 combination therapy in HIV-1_ADA_-infected humanized mice. Graphs display the absolute HIV-1 RNA plasma copies/mL (top) and relative log_10_ changes from baseline viral loads (bottom) after initiation of bNAb therapy. Dashed lines (top graphs) indicate the lower limit of quantitation of the qPCR assay (LLQ) (260 copies/mL). Red lines display the average log_10_ changes compared to baseline viral loads (day −2). **e**, Analyses of single HIV-1 plasma env sequences from HIV-1_ADA_-infected humanized mice obtained after viral rebound for 007 and 10-1074 combination therapy groups. Total number of analyzed sequences is indicated in the center of each pie chart. Mice are labeled according to icon legends in A. Colored bars on the outside of the pie charts indicate mutations in V1/V2 loop and V3 loop. Sensitivity (IC_50_s) of pseudoviruses generated from SGS-derived sequences against 007 and 10-1074. **f**, Alignment of plasma SGS-derived *env* sequences obtained from individual mice (y-axis) after viral rebound following mono- or sequential therapy. *Env* sequences are shown as horizontal gray bars from residues 100-500 relative to HXB2 (x-axis). Mutations identified from day −2 are indicated in black, mutations after viral rebound following monotherapy in red (panels **a** and **b**), and following administration of 007 and 10-1074, either sequentially or in combination in blue (panels **c**, **d** and **e**). Sites at which selected mutations can confer resistance are highlighted by vertical blue bars.

The distinct viral escape profile of 007 led us to explore whether dual-targeting the conserved V3 glycan site could force the virus to accumulate mutations in this conserved epitope, potentially reducing viral fitness and prolonging viral suppression. To investigate this, we sequentially administered 007 IgG to animals pretreated with 10-1074 (and vice versa) once their viral loads rebounded to baseline levels due to the emergence of resistant escape variants (1 mg loading dose followed by 0.5 mg twice weekly for each antibody) (Fig. 6c). To maintain selection pressure, the initial antibody therapy (10-1074 or 007) was continued while the complementary antibody was added. Despite circulating viral variants resistant to the initially applied antibody, sequential addition of 007 to 10-1074 treatment (and vice versa) led to a transient suppression of viremia in all treated animals, indicating that 007 can overcome 10-1074 escape *in vivo* and vice versa (Fig. 6c). Next, we examined whether initiating treatment with a combination of 007 and 10-1074 IgGs would exert greater selection pressure and demonstrate enhanced *in vivo* efficacy compared to monotherapy or sequential combination therapy (Fig. 6d). Administration of combination therapy resulted in prolonged suppression of viremia with viral rebound occurring 42 days after treatment initiation demonstrating *in vivo* synergy (Fig. 6d). SGS of plasma rebound viruses from the sequential and initial combination therapy group revealed selection of combined escape mutations in the V1/V2 and V3 loop residues previously identified from the antibody monotherapy groups (N332_gp120_, 146_gp120_, 303_gp120_ or 322_gp120_) (Fig. 6e,f, Supplementary Table 11). Our findings demonstrate that antibody 007 can overcome 10-1074-class viral escape *in vivo* and that a V3 dual-targeting strategy enhances *in vivo* efficacy.

## Discussion

Advances in donor screening and antibody isolation techniques have accelerated the discovery of potent, broadly neutralizing anti-HIV-1 antibodies^24,25,41,42,48,85–88^. Clinical trials have demonstrated their promise for prevention and treatment of HIV-1 infections and informed vaccine design efforts^4–7,15,16,89^. However, similar to monotherapy with antiretroviral agents, administration of single bNAbs results in the rapid selection of resistant viral variants and fails to achieve the requisite antiviral activity for clinical success^11,46,76,90–93^. Application of bNAb combinations with complementary neutralization coverage have demonstrated enhanced neutralizing activity and long-term control of viremia in preclinical and clinical settings^10,12,13,44,45,94,95^. Similarly, the development of a fully protective vaccine will likely require the induction of bNAbs targeting multiple HIV-1 Env epitopes, emphasizing the need for the identification and characterization of bNAbs directed against novel antigenic sites.

In this study, we characterized the new anti-HIV-1 bNAb 007 targeting the V3 glycan site of the Env trimer through a distinct binding mode. The V3 glycan site is centered around the PNGS at residue N332_gp120_ and extends to high mannose and complex-type N-glycans at positions N156_gp120_, N295_gp120_, N301_gp120_, N339_gp120_, N386_gp120_, and N392_gp120_ as well as the underlying protein surface^21^. bNAb lineages directed against the V3 glycan site recognize the ^324^GD/NIR^327^ protein motif, exhibiting variability in N-glycan accommodation and binding angles of approach^24,25,28,63^. Unlike classical V3 glycan site bNAbs^24,25,76^, 007 does not require the N332_gp120_ glycan for binding. Instead, it interacts primarily with glycans at positions N156_gp120_ and N301_gp120_. In terms of glycan dependency, 007 shares similarities with the recently identified NAb EPTC112^30^. However, it differs in its CDRH3-mediated mode of binding, more pronounced interactions with the ^324^GD/NIR^327^ protein motif, and, most importantly, its superior antiviral activity. While EPTC112 neutralized only 26% (cut-off ≤10µg/mL, 110 strains) of HIV-1 strains from the 119 multiclade panel with low potency^31^, 007 ranks among the most potent and broad V3 glycan site bNAbs identified to date. The enhanced neutralizing activity of 007 is an important consideration for the potential utility of this epitope for vaccine design, therapy, and/or prevention.

The V3 glycan site represents a prime target for vaccine design^89,96–102^ as bNAb responses directed against the N332_gp120_-supersite are among the most frequently elicited in HIV-1-infected individuals^103,104^. Moreover, the diverse binding poses, varying glycan dependencies, distinct antibody gene usage, and the moderate level of somatic hypermutation observed in some V3-targeting bNAbs further underscore the potential of the V3 glycan site as a favorable target for immunogen design^22,63,105^. Whereas current immunization strategies seek to elicit N332_gp120_ glycan-specific antibody responses^89,96–102^, the N332_gp120_-independent V3 epitope has not been considered. In this context, 007 represents a promising avenue for vaccine development due to its enhanced antiviral activity and sequence features. Similar to the V3 bNAb BG18, a target of vaccine design^89,97^, 007 exhibits a shorter CDRH3 compared to other characterized V3-targeting bNAbs and lacks insertions or deletions in both its heavy and light chains^89,97,106,107^. These features may enhance the feasibility of eliciting 007-like bNAbs by vaccination. However, the high degree of somatic hypermutation in 007 could pose a challenge for inducing similar bNAbs via vaccination, necessitating germline reversion studies to determine whether minimally mutated variants retain antiviral activity^97,108^.

Due to its distinct epitope and neutralization profile, 007 represents a promising potential candidate for therapy, functional cure, and prevention strategies involving passive administration of bNAb combinations. This is of particular interest in regions of the world with a high prevalence of CRF01 AE viruses that represent a coverage gap of V3 glycan site bNAbs due to the lack of the N332_gp120_ glycan^109^. Indeed, 007 retains high levels of antiviral activity against CRF AE and AC viruses and could efficiently substitute classical V3 glycan site bNAbs in combination regimens considered for these regions. The distinct V3-binding mode and escape profile of 007 have important clinical implications, as they contribute to 007’s potential to effectively complement other V3 glycan site bNAbs, which, despite their high potency, often exhibit limited breadth^24,25,28^. The ability of 007 to overcome 10-1074 escape both *in vitro* and *in vivo*, along with its predicted combined breadth of 86.4% in combination with 10-1074, highlight new opportunities for bNAb combination strategies, including dual targeting of the V3 glycan site to enhance breadth, potency and selection pressure on this conserved epitope.

Structural analysis of the 007 Fab in complex with a SOSIP Env trimer revealed sub-stoichiometric Fab binding, a surprising finding for a potent bNAb. Interestingly, EPTC112, which recognizes a similar epitope, also lacked full Fab occupancy in its structure, with only two Fabs per trimer modeled^30^. Similar to results with EPTC112^30^, *in vitro* neutralization assays for 007 demonstrated IgG bivalency enhanced neutralization potencies. Based on our 007 Fab-SOSIP structures, we hypothesized that intra-spike bivalent IgG binding could occur on open forms of an Env trimer. However, when SOSIP-IgG complexes were analyzed by cryo-EM, a trimer-dimer of closed Env-IgG complexes was observed, a configuration that could occur if IgGs link Envs trimers on two separate virions. Analogous structures linking two SARS-CoV-2 spike trimers by IgGs were postulated to enhance neutralization by aggregating virions^32^, and cryo-electron tomography of nAbs incubated with SARS-CoV-2 demonstrated inter-virion bivalent binding of IgGs^110^. It should be noted that the SOSIP Env trimers we used for structure determinations contained specific mutations (I559P_gp41_, A501C_gp120_, T605C_gp41_) that decrease the sampling of alternative Env conformations^111^, whereas neutralization assays use membrane-bound Env trimers lacking these mutations. Further studies of 007 IgG in complex with HIV-1 virions or virus-like particles displaying native Env trimers from strains with high MNRs could elucidate its mechanism of bivalent neutralization. In summary, the favorable neutralization properties and distinct viral escape profile position 007 as a promising investigational candidate for HIV-1 therapy, functional cure, and prevention. Our findings further emphasize the N332_gp120_ glycan-independent V3 epitope as a compelling target for vaccine development.

## Methods

### Study participants and collection of clinical samples

Large blood draws and leukapheresis samples were collected in accordance with protocols reviewed and approved by the Institutional Review Board (IRB) of the University of Cologne (study protocols 13-364 and 16-054) and local IRBs. Study participants were recruited from private practices and/or hospitals in Germany (Cologne, Essen, and Frankfurt), Cameroon (Yaoundé), Nepal (Kathmandu), and Tanzania (Mbeya), with all participants providing written informed consent. A total of 2,392 serum samples were screened for anti-HIV-1 neutralizing activity to identify HIV-1 elite neutralizers^35^. Study individual EN01 was selected for a large blood draw and subsequent B cell isolation.

### Cell lines

HEK293T cells were cultured in Dulbecco’s Modified Eagle Medium (DMEM, Thermo Fisher), supplemented with 10% fetal bovine serum (FBS, Sigma-Aldrich), 1x antibiotic-antimycotic (Thermo Fisher), 1 mM sodium pyruvate (Gibco), and 2 mM L-glutamine (Gibco) at 37°C in an atmosphere containing 5% CO_2_. HEK293-6E cells were grown in FreeStyle 293 Expression Medium (Life Technologies), supplemented with 0.2% penicillin/streptomycin, and maintained under constant agitation at 90–120 rpm at 37°C and 6% CO2. TZM-bl cells (Platt et al., 1998) were cultured in DMEM supplemented with 10% FBS, 1 mM sodium pyruvate, 2 mM L-glutamine, 50 µg/mL gentamicin (Merck), and 25 mM HEPES (Millipore) at 37°C in 5% CO2. All three cell lines (HEK293T, HEK293-6E, and TZM-bl) were of female origin and were not specifically authenticated.

### Mouse Models

NOD.Cg-Rag^1tm1mom^Il2rg^tm1Wjl^/SzJ (NRG) mice were acquired from The Jackson Laboratory and subsequently bred and housed within the Decentralized Animal Husbandry Network (Dezentrales Tierhaltungsnetzwerk) at the University of Cologne. Mice were maintained under specific pathogen-free (SPF) conditions with a 12-hour light/dark cycle. Breeding mice were provided ssniff 1124 breeding feed, while experimental mice received ssniff 1543 maintenance feed. The generation of humanized mice followed an established protocol with slight modifications^45,113^. Human CD34⁺ hematopoietic stem cells were isolated from umbilical cord blood and placental tissue through immunomagnetic separation using CD34 microbeads (Miltenyi Biotec). The collection of these tissue sources was conducted with prior written informed consent, following protocols approved by the Institutional Review Board of the University of Cologne (16-110) and the Ethics Committee of the Medical Association of North Rhine (2018382). Within five days after birth, NRG mice received sublethal irradiation, after which human CD34⁺ stem cells were administered via intrahepatic injection 4 to 6 hours later. The engraftment and humanization efficiency were assessed 12 weeks post-injection using FACS to detect PBMCs in circulation^45^. All animal experiments were performed in compliance with ethical regulations and approved by the State Agency for Nature, Environmental Protection, and Consumer Protection of North Rhine-Westphalia (LANUV).

### PBMCs and plasma isolation

Peripheral blood mononuclear cells (PBMCs) were isolated from large-volume blood samples using density gradient centrifugation with Histopaque separation medium (Sigma-Aldrich) and Leucosep cell tubes (Greiner Bio-One), following the manufacturer’s instructions. The purified PBMCs were cryopreserved at −150°C in a freezing medium composed of 90% fetal bovine serum (FBS) and 10% dimethyl sulfoxide (DMSO) until further use. Plasma samples were collected separately and stored at −80°C for subsequent analyses.

### Isolation of single HIV-1-reactive B cells

The isolation of single antigen-reactive B cells was carried out following previously established methods^39,114^. CD19⁺ B cells were selectively enriched from PBMCs using immunomagnetic separation with CD19 microbeads (Miltenyi Biotec) according to the manufacturer’s protocol. The isolated CD19⁺ B cells were subsequently stained on ice for 20 minutes with 4’,6-diamidino-2-phenylindole (DAPI; Thermo Fisher Scientific), anti-human CD20-Alexa Fluor 700 (BD), anti-human IgG-APC (BD), and GFP-labeled BG505_SOSIP.664_^37^. Following staining, DAPI⁻ CD20⁺ HIV-1 Env-reactive IgG⁺ single cells were sorted into 96-well plates using a FACSAria Fusion cell sorter (Becton Dickinson). Each well was preloaded with 4 µl of sorting buffer containing 0.5× PBS, 0.5 U/µl RNAsin (Promega), 0.5 U/µl RNaseOut (Thermo Fisher Scientific), and 10 mM dithiothreitol (DTT; Thermo Fisher Scientific). The plates were immediately cryopreserved at −80°C following cell sorting.

### Amplification and analysis of heavy and light chain V gene sequences

Antibody heavy and light chain amplification from single cells was primarily conducted as described in prior studies^39,41,115^. Reverse transcription was carried out using Random Hexamers (Invitrogen) and Superscript IV (Thermo Fisher Scientific) in the presence of RNase inhibitors RNaseOUT (Thermo Fisher Scientific) and RNasin (Promega) to preserve RNA integrity. The synthesized cDNA was subsequently employed for the amplification of immunoglobulin heavy and light chains using PlatinumTaq HotStart polymerase (Thermo Fisher Scientific), supplemented with 6% KB extender and gene-specific primer mixes targeting V gene regions. A semi-nested PCR strategy was applied with optimized V gene-specific primer mixes^38^ to improve amplification efficiency^115,41,38,39^. The resulting PCR products were evaluated via gel electrophoresis to confirm expected fragment sizes before undergoing Sanger sequencing Raw sequence processing, annotation and clonal assignment was performed with the Antibody Repertoire Toolkit (AbRAT)^116^ using default settings. In brief, chromatograms were filtered to only retain sequences with a mean Phred quality score of at least 28 and a minimum read length of 240 nucleotides (nt). Variable region annotation, spanning from FWR1 to the end of the J gene segment, were annotated based on IgBLAST^117^. Nucleotide positions within the variable region exhibiting Phred scores below 16 were masked, and sequences containing more than 15 masked bases, premature stop codons, or frameshifts were excluded from downstream analyses.

For clonal lineage assignment, productive heavy chain sequences were grouped by identical V_H_/J_H_ gene usage and clustered with AbRATs “Iterative Greedy CDR3 Clustering”-algorithm that is based on the pairwise Levenshtein distance between CDRH3s. Clonal clusters were defined by initiating the grouping process from a randomly selected sequence, with membership requiring a minimum of 75% CDRH3 amino acid identity (relative to the shortest sequence). To enhance classification accuracy, 100 iterations of randomization and clonal assignment were performed, with the configuration yielding the lowest number of unassigned sequences chosen for subsequent analysis. All assigned clone groups were manually validated by investigators, incorporating shared somatic mutations and light chain pairing information to ensure consistency in lineage identification.

### Cloning and production of monoclonal antibodies

The heavy and light chain V gene regions of mAbs were synthesized as eBlocks gene fragments (IDT), incorporating complementary overhangs pre-configured for cloning into expression vectors (IgG1, Igλ, Igκ). Cloning was performed using Sequence- and Ligation-Independent Cloning (SLIC) with T4 DNA polymerase (New England Biolabs) and chemically competent *Escherichia coli* DH5α, following established protocols^118,41,115,39,114^. Positive transformants were identified through Sanger sequencing, after which confirmed bacterial colonies were expanded in LB medium. Plasmid DNA was subsequently extracted using the NucleoBond Xtra Midi kit (Macherey-Nagel).

MAbs were generated by co-transfecting HEK293-6E cells (0.8 × 10⁶ cells in 50 mL) with heavy chain (IgG1) and light chain (Igλ or Igκ) expression plasmids, employing polyethylenimine (PEI; Sigma-Aldrich) as the transfection reagent. Cells were maintained at 37°C with 5% CO₂ in FreeStyle 293 Expression Medium (Thermo Fisher Scientific), supplemented with 0.2% penicillin/streptomycin, under continuous agitation at 120 rpm. Culture supernatants were harvested seven days post-transfection via centrifugation and incubated overnight at 4°C with Protein G-coupled beads (GE Life Sciences). The beads were subsequently transferred to chromatography columns (Bio-Rad), washed with DPBS (Thermo Fisher Scientific), and antibodies were eluted using 0.1 M glycine (pH 3) into 1 M Tris (pH 8) to neutralize acidity. A final buffer exchange to PBS was performed using 30K Amicon spin membranes (Merck Millipore). Antibody concentrations were determined via UV spectrophotometry using a Nanodrop system (Thermo Fisher Scientific).

007 and 10-1074 IgGs used for structural studies, mass spectrometry, and molar neutralization ratio assays were expressed via transient co-transfection of Expi293F cells with IgG1 heavy and light chain plasmids. IgGs were purified from cell culture supernatant using MabSelect™ SuRe (Cytiva), concentrated, and SEC purified on a Superdex 200 16/60 column (Cytiva). SEC fractions corresponding to IgG were combined and concentrated.

To produce Fabs, the heavy chain variable region of 007 was subcloned into a mammalian expression vector containing the CH1 domain and a C-terminal 6xHis tag. 10-1074 plasmids were cloned as previously described^55^. Fab heavy and light chain plasmids were used to transiently co-transfect Expi293F cells (Thermo Fisher) and Fabs were purified from culture supernatant by immobilized metal affinity chromatography (IMAC) using a HisTrap HP column (Cytiva). Fabs were concentrated and buffer exchanged into TBS (20mM Tris pH 8.0, 150mM NaCl) using Amicon 10 kDa spin concentrators (Millipore) and further purified by size exclusion chromatography (SEC) on a Superdex 200 16/60 column (Cytiva) equilibrated with TBS. SEC fractions corresponding to Fab were combined and concentrated.

To produce bispecific IgGs comprising a 007 arm and an anti-CD3 OKT3 arm^65^, mutations were introduced into the respective IgG1-LS^119^ constant regions (E357Q and S364K in 007_HC_ and Q295E, L368D, K370S, N384D, Q418E, N421D in OKT3_HC_) to promote heavy chain heterodimerization and facilitate downstream purification^64^, and the CH1 and CL domains of the OKT3 antibody were domain swapped to promote correct heavy and light chain pairing of both Fab arms^120^. Heavy and light chain plasmids were mixed in equal amounts (50 µg of each of the four plasmids per 200mL transfection culture) and used to transiently co-transfect Expi293F cells (Thermo Fisher). IgGs were purified from cell culture supernatant using a MabSelect™ SuRe (Cytiva) affinity column and were concentrated and buffer exchanged into 50mM Tris, pH 8.7 using 10 kDa Amicon spin concentrators (Millipore). Bispecific IgGs were purified by anion exchange chromatography using a HiTrap Q column (Cytiva) and eluted with a NaCl gradient. Fractions corresponding to the bispecific IgG were combined and concentrated before SEC purification using a Superdex 200 16/60 column (Cytiva) equilibrated in TBS.

### Expression and purification of BG505 SOSIP trimers

BG505^52^ and BG505-DS^53,54^ SOSIPs used for cryo-EM and mass spectrometry experiments were expressed via transient co-transfection of Expi293F cells with a plasmid encoding soluble furin. Briefly, SOSIPs were purified from cell culture supernatant by PGT145 (BG505) or 2G12 (BG505-DS) immunoaffinity chromatography, dialyzed in TBS, concentrated to <2 mL, and purified by SEC on a Superose 6 Increase column (Cytiva) (BG505) or sequentially on a Superdex 200 16/60 column (Cytiva) followed by Superose 6 Increase column. SEC fractions corresponding to trimeric SOSIPs were combined and concentrated.

### Quantification of unpurified mAbs from cell supernatants by human IgG capture ELISA

A human IgG capture ELISA was employed to measure antibody concentrations in unpurified supernatants from transfected HEK293-6E cells, with slight modifications to established protocols^39^. ELISA plates (Greiner Bio-One) were coated with 2.5 µg/mL polyclonal goat anti-human IgG (Jackson ImmunoResearch) in PBS and incubated for at least 45 minutes at 37°C or alternatively overnight at 4°C. Following the coating step, plates were blocked for 60 minutes at room temperature (RT) with a blocking buffer (BB) composed of PBS supplemented with 5% nonfat dry milk powder (Carl Roth T145.2). Supernatants from transfected HEK293-6E cells were diluted 1:20 in BB before analysis, while a human myeloma IgG1 kappa standard (Sigma-Aldrich) was prepared at an initial concentration of 4 µg/mL in BB. Both samples and standards were subjected to serial 1:3 dilutions in BB and incubated for 45 minutes at RT. Detection was performed using an HRP-conjugated anti-human IgG antibody (Southern Biotech 2040-05) diluted 1:2500 in BB. Colorimetric development was initiated by the addition of ABTS substrate (Thermo Fisher Scientific 002024), and absorbance was measured at 415 nm and 695 nm using a microplate reader (Tecan). Antibody concentrations in the supernatants were calculated by interpolation from the human IgG1 standard curve.

### Generation of HIV-1 pseudoviruses

HIV-1 pseudoviruses were produced in HEK293T cells by co-transfection with pSG3Δenv and the respective HIV-1 Env plasmids as previously described^1,2,36,84^. A synthetic HIV-1_ADA_ Env plasmid was obtained from Twist Bioscience for the production of HIV-1_ADA_ pseudoviruses containing point mutations identified through *in vivo* single-genome sequencing (SGS) analysis. Site-specific mutations were introduced using the Q5 Site-Directed Mutagenesis Kit (New England Biolabs) according to the manufacturer’s instructions. Kifunensine-treated pseudoviruses were produced under presence of 5 µg/ml Kifunensine. Supernatants containing Kifunensine-treated pseudoviruses were harvested 3 days after transfection and stored at −80°C.

### Generation of mutant HIV-1 pseudoviruses

Mutant variants of HIV-1 pseudoviruses were generated by introducing site-specific mutations into gp160 expression plasmids. Point mutations were incorporated using the Q5 Site-Directed Mutagenesis Kit (New England Biolabs) following the manufacturer’s protocol. The resulting mutant plasmids were subsequently used for pseudovirus production, following the protocol as described above for wild-type pseudoviruses.

### Determination of neutralizing activity by luciferase-based TZM-bl assays

Neutralization assays were performed to determine the IC₅₀ and IC₈₀ values of purified mAbs and to assess neutralizing activity in unpurified HEK293-6E cell culture supernatants. These assays were performed with slight modifications to previously described protocols^84,121,122^. Purified mAbs, serum IgGs, or HEK293-6E cell culture supernatants were pre-incubated with HIV-1 pseudovirus strains for 1 hour at 37°C before adding 10⁴ TZM-bl cells per well in a 96-well plate. After 48 hours of incubation at 37°C with 5% CO₂, luciferase activity was measured using a luciferin/lysis buffer. Background RLUs from non-infected control wells were subtracted, and the percentage of neutralization was calculated. For screening unpurified IgGs in HEK293-6E cell supernatants, a final IgG concentration of 2.5 µg/mL was used, as determined by human IgG capture ELISA. For large-scale donor screening, IgGs isolated from participant samples were tested against each pseudovirus at a fixed concentration of 300 µg/mL in duplicate wells. IC₅₀ and IC₈₀ values for mAbs were determined by performing serial dilutions starting at 10, 25, or 50 µg/mL. These values, representing the mAb concentration required to reduce viral signal by 50% or 80%, were calculated using a dose-response curve fitted in GraphPad Prism. All IC₅₀ and IC₈₀ determinations were performed in duplicates for each mAb.

### Molar neutralization ratio (MNR) assays

Pseudovirus neutralization assays were conducted using TZM-bl reporter cells as described^121,122^. IgGs, bispecifics, and Fabs were evaluated in duplicate with an eight-point, five-fold dilution series starting at a top concentration of 50 μg/mL (007 bispecific, 007 Fab, 10-1074 IgG, 10-1074 Fab) or 2 μg/mL (007 IgG). The dilution at which 50% of virus was neutralized (IC_50_) is reported in μg/mL and molar concentrations in Supplementary Table 6. MNRs were calculated as the ratio of molar IC_50_ values for different formats of the antibody.

### Determination of antibody interference by competition ELISAs

To assess antibody interference, monoclonal antibodies were biotinylated using the EZ-Link Sulfo-NHS-Biotin Kit (Thermo Fisher Scientific) according to the manufacturer’s protocol. Excess biotin was removed by buffer exchange into PBS using Amicon 10 kDa centrifugal filter membranes (Millipore). High-binding ELISA plates (Greiner Bio-One) were coated overnight at 4°C with an anti-6×His tag antibody (Abcam 9108) at a final concentration of 2 µg/mL. After coating, plates were blocked for 1 hour at 37°C with PBS supplemented with 3% bovine serum albumin (BSA; Sigma-Aldrich) to prevent nonspecific binding. BG505_SOSIP.664_-His protein was then added at 2 µg/mL in PBS and incubated for 1 hour at room temperature (RT) to facilitate antigen capture. To evaluate competition, unlabeled antibodies were applied at an initial concentration of 32 µg/mL in PBS and subjected to a 1:3 serial dilution. After a 1-hour incubation at RT, biotinylated mAbs were introduced at 0.5 µg/mL in PBS containing 3% BSA, followed by another 1-hour incubation at RT. Detection was performed using peroxidase-conjugated streptavidin (Jackson ImmunoResearch) diluted 1:5,000 in PBS containing 1% BSA and 0.05% Tween-20. Between each incubation step, wells were thoroughly washed with PBS containing 0.05% Tween-20 (Carl Roth) to remove unbound components. Colorimetric detection was achieved using ABTS substrate solution (Thermo Fisher Scientific 002024), and absorbance was measured at 415 nm and 695 nm using a microplate reader (Tecan).

### Neutralization fingerprint analyses

The neutralizing activity against the f61 pseudovirus^36^ panel was analyzed by calculating Spearman correlation coefficients for each pair of antibodies based on their neutralization data, and visualizing the results as a heatmap (Fig. 1d). Neutralization fingerprint analysis (Fig. 1e) was performed based on a diverse panel of 24 HIV-1 strains for which IC_50_s were available for all antibodies using an approach described in^48^. An antibody-antibody distance matrix was calculated pairwise as the sum over the panel of the absolute differences of the log IC_50_s. The dendrogram was calculated from this distance matrix using hierarchical clustering by the R command “hclust” using the “average” method.

### Assessment of autoreactivity in HEp-2 cell assays

Autoreactivity of mAbs was evaluated using the NOVA Lite HEp-2 ANA Kit (Inova Diagnostics) following the manufacturer’s protocol. Antibodies were applied at a final concentration of 100 µg/mL in PBS to HEp-2 cell-coated slides. After incubation and subsequent washing steps, fluorescence imaging was performed using a DMI 6000 B fluorescence microscope (Leica) under standardized conditions: a 3-second exposure time, 100% light intensity, and a gain setting of 10. Fluorescent signals were analyzed to determine autoreactivity profiles.

### Cryo-EM sample preparation

007 Fab was incubated in a 3.4:1 molar ratio of Fab to BG505-DS SOSIP trimer^53,54^ and incubated overnight at room temperature. SOSIP trimers and Fab-SOSIP complexes were purified from unbound Fabs on a Superose 6 10/300 Increase column (Cytiva) operating in TBS and concentrated using a 30 kDa spin concentrator to ∼3.4 mg/mL immediately prior to vitrification. 007 IgG1 was added to BG505 SOSIP at a 1:1 molar ratio of IgG to BG505 SOSIP trimer, with a total protein concentration of ∼2.6 mg/mL and vitrified after incubating for ∼38 hours at room temperature.

Octyl-maltoside, fluorinated solution (Anatrace) was added to 0.02% (w/v) final concentration for each sample immediately before addition of 3 µL to a Quantifoil R1.2/1.3 Cu 300 mesh grid (Electron Microscopy Sciences) that had been glow discharged for 1 min at 20 mA using a PELCO easiGlow™ (Ted Pella). Grids were blotted for 3-4s with Whatman No. 1 filter paper and vitrified in liquid ethane using a Mark IV Vitrobot (Thermo Fisher) operating at 22°C and 100% humidity.

### Cryo-EM data collection and processing

Data for the 007 Fab-SOSIP sample were collected on a 300 keV Titan Krios transmission electron microscope (Thermo Fisher) equipped with a Gatan BioQuantum Energy Filter and a K3 6k x 4k direct electron detector, and data for the 007 IgG1-SOSIP sample were collected on a 200 keV Talos Arctica (Thermo Fisher) equipped with a Gatan K3 6k x 4k direct electron detector. 40-frame movies were collected in SerialEM^123^. 007 Fab-SOSIP movies were recorded in super-resolution (0.416 Å per pixel) using a 3×3 beam image shift pattern with 3 shots per hole and 007 IgG-SOSIP movies were recorded in super-resolution (0.72 Å per pixel) with a 3×3 beam image shift pattern and 1 shot per hole.

Data collection and processing details are included in Supplementary Table 4 and Extended Data Fig. 4, 8. Motion correction, CTF estimation, particle picking, and particle extraction were performed in cryoSPARC Live v4^124^. Extracted particles were subject to 2D classification in cryoSPARC^124^ and particles from select 2D classes processed by 3D classification in RELION v4.0.1^125,126^. Particles from select 3D classes were re-extracted in cryoSPARC and subject to ab initio reconstruction and non-uniform refinement^124,127^. Particles underwent reference-based motion correction and were subject to a final round of non-uniform refinements^127^.

### Model Building and Refinement

Initial coordinates for the 1 Fab-bound trimer structure were generated by docking individual protein chains from reference structures [PDB 5BZD (V_H_), 7PS3 (V_L_), and 6UDJ (BG505)] into the corresponding EM density in ChimeraX^128,129^. The initial model was sequence corrected in Coot^130^ and underwent iterative rounds of refinement in Phenix^131^ and Coot^130^. Glycans were built in Coot^132^ and glycan geometries evaluated in Privateer^133^. Coordinates for the 1 Fab-bound trimer aided in generating trimer structures with 0, 2, or 3-bound Fabs, as well as the 3 IgG-bound trimer-dimer structure. To facilitate measurements between the C-termini of the Fab heavy chains, the Fab C_H_C_L_ domains from PDB 8UKI were docked into the corresponding densities in the 2 Fab-bound trimer structure and the 3 IgG-bound trimer-dimer structure. Antibodies were numbered according to Kabat.

### Structural analyses

Figures were prepared using UCSF ChimeraX v1.9^128,129^. 007 Fab-Env interactions were evaluated using the 1 Fab-bound trimer structure. Distance measurements between C-termini of Fab HCs were taken from the alpha carbon of R210 in the CH1 domain (analogous to residue 222) in either the 2 Fab-bound trimer structure or the 3 IgG-bound trimer-dimer structure. Coordinates for a 007-bound gp120 were aligned to two gp120 chains in a b12-bound trimer structure (PDB 5VN8)^134^ to model the analogous distance on an occluded open conformation of the trimer. The distance between trimer apexes reported for the trimer-dimer structure was measured between residues N188 on opposing trimers.

### Immunoprecipitation for site-specific N-glycan analyses

To isolate BG505 trimers recognized by 007_IgG_ (007_IgG_-bound BG505), we developed an optimized protocol for immunoprecipitation. Briefly, BG505 and 007_IgG_ were mixed at a 1:2 (w:w) ratio and incubated overnight at 4°C in 20 mM sodium phosphate buffer (pH 7.2). 007_IgG_-bound BG505 complexes were isolated using affinity chromatography (Protein G Sepharose; GE Healthcare Life sciences Corp., Piscataway, NJ). Protein G Sepharose was equilibrated with 20 mM sodium phosphate buffer (pH 7.2) and incubated overnight at 4°C with the preincubated mixture of BG505 and 007_IgG_. 007_IgG_-BG505 complexes captured by Protein G Sepharose were separated from unbound BG505 by centrifugation at 1,000 x g for 3 min, eluted with 0.1 M glycine (pH 2.5), and neutralized with 1 M Tris-HCl (pH 9.0). Samples were then stored at −20°C.

### Enzymatic removal of N-glycans from Env BG505 SOSIP

Total BG505 and 007_IgG_-bound BG505 samples were denatured for 10 min at 100°C in a denaturing buffer provided with peptide N-glycosidase F (PNGase F, Prozyme, Hayward, CA). After samples were chilled on ice for 5 min, PNGase F was added together with detergent following the manufacturer’s instructions (PNGase F, Prozyme, Hayward, CA). Samples were then incubated at 37°C for 30 h.

### Isolation of gp120 component chains of BG505 SOSIP for LC-MS

Total BG505 and 007_IgG_-bound BG505 samples were separated by SDS-PAGE under reducing conditions on 10% Mini-PROTEAN TGX precast gels (Bio-Rad Laboratory Inc.). Gels were briefly washed with water and stained with Bio-safe colloidal Coomassie G-250 Stain (Bio-Rad Laboratories, Inc., Hercules, CA). After destaining, protein bands corresponding to natively glycosylated gp120 (∼130 kDa) or samples deglycosylated with PNGase F (∼65 kDa) were excised from the gel and stored at −20°C for LC-MS analyses (Extended Data Fig. 6).

### LC-MS and MS/MS analysis of gp120

The excised bands were digested with trypsin (Promega) and extracted from the gel matrix by use of standard in-gel protease digestion methods^135,136^. The resulting peptide/glycopeptide mixtures were analytically separated on a self-prepared C18 reversed-phase pulled-tip column using a nano-liquid chromatography (nano-LC) system as previously described^135,136^. The eluted glycopeptides were electrosprayed at 2 kV into a dual linear quadrupole ion trap Orbitrap Velos Pro mass spectrometer (Thermo Fisher, San Jose, CA). The mass spectrometer was set to switch between a full scan (400 < m/z < 2,000) followed by successive MS/MS (200 < m/z < 2,000) scans of the 10 most abundant precursor ions using the collision-induced dissociation (CID) method.

### Glycopeptide identification and quantitation

Site-specific N-glycan heterogeneity profiles of gp120 from total BG505 and 007_IgG_-bound BG505 samples were determined using a workflow similar to before^135,136^, which consisted of three main steps:

*Step 1*. *Initial glycopeptide identification*: LC-MS/MS data for the deglycosylated gp120 were analyzed using the Single Protein Screening and Quantitation workflow in the Pinnacle software (version 1.0.103 Optys Tech Corporation). Identification of peptides containing specific N-glycosylation sites (NGSs) was achieved with a peptide tolerance of 10 ppm in MS1 and an MS/MS tolerance of 0.7 Da. All peptide assignments were validated by visual inspection of the associated MS1 and MS/MS spectra. An increase of 1 Da in the peptide mass value of the deglycosylated gp120 indicated the presence of N-glycosylation site with an attached glycan in the intact gp120 because PNGase F treatment converted each glycosylated Asn into Asp. Site occupancy was calculated based on the areas under the curve (AUC) for the peptide containing the unmodified Asn and the peptide containing the Asp.

*Step 2. Glycopeptide quantitation:* To identify the full range of glycan heterogeneity at specific sites, LC-MS/MS data for the intact gp120 were analyzed using the Targeted Quantitation – Label Free DDA workflow in the Pinnacle software. The search was conducted with a 10-ppm mass accuracy of the 3 most abundant isotopes with a series of custom peptide and glycopeptide workbooks generated for each NGS. To speed up the validation process and reduce the number of false-positive results, the confirmed deglycosylated peptides from *Step 1* were used to limit the retention time (RT) window search for each glycopeptide. Once the glycopeptides were identified, the AUC from each glycopeptide was expressed as a percent relative abundance of the total sum of all glycoforms for an NGS.

*Step 3. Visualization of N-glycan heterogeneity at specific sites*: The entire range of glycans at a single site was presented as a stacked bar divided into the relative distributions of the broad N-glycan categories of high-mannose, hybrid, and complex, using the same color scheme as previously described^135^. To better visualize changes in glycoforms between total BG505 and 007_IgG_-bound BG505, a side-by-side bar graph was also included which differentiated each high-mannose glycan but kept the hybrid and complex as single groups. Error bars represent the standard deviations of replicate measurements (three for total BG505 and two for 007_IgG_-bound BG505). Differences between groups were evaluated for statistical significance based on P-values calculated using Welch’s T-test.

### Deep mutational scanning

A previously established lentiviral deep mutational scanning platform was utilized to assess the effects of all mutations on HIV Env escape from antibodies^77,79^. This system employs barcoded pseudoviruses to facilitate comprehensive mutational analysis. The BF520 Env and TRO.11 Env mutant libraries were generated in prior studies^1,3^ and were reused in the present investigation. Detailed descriptions of the construction and composition of these mutant libraries are available in previous reports^77,79^. TRO.11 Env function of entry into cells and escape from antibody 10-1074 are published in these prior studies^77,79^. In this study, the BF520 and TRO.11 Env mutant libraries were employed to map escape from antibodies 007, BG18, PGT121, and PGT128. For each mutant library, a small proportion of VSV-G pseudotyped viruses carrying unique barcodes was introduced as an internal infection control, as these viruses are not susceptible to HIV-1-specific antibodies. Each library was incubated for one hour with antibodies at concentrations ranging from IC₉₀ to IC₉₉.₉, alongside a control incubation without antibody. TZM-bl cells were then infected with the antibody-treated pseudovirus libraries, and 12 hours post-infection, unintegrated lentiviral genomes were extracted using a miniprep approach. Variant barcodes were subsequently amplified and sequenced following previously established protocols^79^.

### Deep mutational scanning data analysis

Deep mutational scanning data were analyzed using *dms-vep-pipeline-3* version 3.20.1. See https://github.com/dms-vep/HIV_Envelope_TRO11_DMS_007 and https://github.com/dms-vep/HIV_Envelope_BF520_DMS_007 for GitHub repositories containing the analyses of the deep mutational scanning data. See https://dms-vep.org/HIV_Envelope_TRO11_DMS_007 and https://dms-vep.org/HIV_Envelope_BF520_DMS_007 for HTML renderings of key analyses, plots, and data files produced by the analyses. Effects of mutations on Env function of entry into cells measured in prior studies^77,79^ were used to shade the logoplots in Fig. 5 and Extended Data Fig. 9a Illumina sequencing data of variant barcodes from antibody selection experiments were processed following previously established methods^78,79^. A comprehensive description of sequencing analyses and modeling of mutation effects is provided in^79^, which should be consulted for methodological details. Briefly, the non-neutralized fraction of each barcoded variant was calculated in each antibody selection by comparing the frequency of each barcoded variant to the frequency of the non-neutralized standard viruses between the antibody incubation and mock incubation conditions. The software package *polyclonal*^137^ version 6.14 was used to model the escape effects of individual mutations for each antibody. In Fig. 5a and Extended Data Fig. 9a the height of each amino acid in each logo plots represents the effect of that individual mutation on escape from that antibody, where taller letters represent more escape. The model of the effects of mutations on escape can also be used to predict the level of neutralization of Env mutants at arbitrary antibody concentrations, which we used to calculate the deep mutational scanning measured log fold change in IC_50_ values in Fig. 5b.

### FcγRIIIa signaling Promega Assay

Signaling through FcγR was assessed using an ADCC reporter bioassay from Promega (Promega G9790 (FcγRIIIa-F), G7010(FcγRIIIa-V)). CHO cells overexpressing HIV JRFL envelope (CHO-HIV JRFL ENV) were used as targets. HIV bnAbs were serially diluted 4-fold and added to target cells, 30 min prior to addition of Jurkat reporter cells. After overnight incubation at 37℃, luminescence was measured using the Bio-Glo-TM Luciferase Assay Reagent according to the manufacturer’s instructions.

### ADCC assay

NK cells were isolated from whole blood using the EasySep™ Direct Human NK Cell Isolation Kit (StemCell Technologies #19665) following the manufacturer’s instructions. CHO-HIV JRFL ENV were incubated with titrated HIV bnAbs prior to addition of effector NK cells at 8:1 (FV NK donor) or 9:1 (FF NK donor) in AIM V Medium (Life Technologies #12055091). Each Ab concentration was tested in duplicate. After 4 hr incubation at 37°C, plate was centrifuged, and supernatants transferred to a new plate containing the LDH-substrate prepared according to manufacturer’s instructions (Roche #11644793001 Cytotoxicity Detection Kit (LDH)). Spontaneous release was determined by incubating target cells with medium alone, and maximal lysis by incubation with 1% Triton X-100. Kinetic Absorbance was measured at 492 and 650 nm with a microplate reader, and cytotoxicity was determined using the following equation:

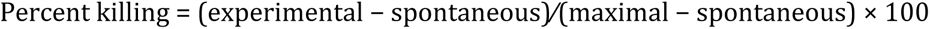

### Generation of replication-competent HIV-1 virus

Recombinant, replication-competent HIV-1_ADA_ variant, incorporating the ADA Env within an NL4-3 backbone, was produced by transfecting HEK293T cells using FuGENE 6 Transfection Reagent (Promega). Virus-containing supernatants were harvested at 48 and 72 h post-transfection, then aliquoted and stored at −80°C for subsequent experiments.

### HIV-1 infection of humanized mice and viral load quantification

Humanized NRG mice were infected intraperitoneally with replication-competent HIV-1_ADA_. At 32 days post-infection, sterile monoclonal antibodies were administered subcutaneously in PBS. A loading dose of 1 mg was initially delivered, followed by maintenance doses of 0.5 mg every 3 to 4 days. To determine viral loads, viral RNA was extracted from EDTA-treated plasma samples using the MinElute Virus Spin Kit (Qiagen), with DNase I treatment (Qiagen) performed on an automated Qiacube system (Qiagen). *pol*-specific primers 5’TAATGGCAGCAATTTCACCA and 5’GAATGCCAAATTCCTGCTTGA, along with the probe 5’/56-FAM/CCCACCAACARGCRGCCTTAACTG/ZenDQ/, as previously described^44^. qRT-PCR was performed using a QuantStudio 5 instrument (Thermo Fisher Scientific) with the TaqMan RNA-to-CT 1-Step Kit (Thermo Fisher Scientific). Heat-inactivated supernatants of replication-competent HIV-1_YU2_ propagated in MOLT-CCR5 cells, were included in each PCR run as a standard for viral load quantification. The viral concentration of this standard was determined using the the cobas 6800 HIV-1 kit (Roche). The quantification limit for qRT-PCR was established at 260 copies/mL. To calculate log_10_ changes in viral load, values below this threshold were assigned a concentration of 260 copies/mL. For normalization, the mean log_10_ viral load change of the PBS control group at each corresponding time point was subtracted from the individual log_10_ viral load changes of the treatment groups.

### Single genome sequencing of plasma HIV-1 *env* from *in vivo* experiments

Single Genome Sequencing (SGS) of HIV-1 *env* genes was conducted following previously established protocols^138^. Briefly, viral RNA was extracted from EDTA-treated plasma samples using the MinElute Virus Spin Kit (Qiagen), followed by DNase I treatment (Qiagen) on an automated Qiacube system (Qiagen). Reverse transcription was performed using the antisense primer YB383 (5’ TTTTTTTTTTTTTTTTTTTTTTTTRAAGCAC) and SuperScript IV reverse transcriptase (Thermo Fisher Scientific) according to the manufacturer’s instructions. To degrade residual RNA, the reaction was treated with 0.25 U/µL RNase H (Thermo Fisher Scientific) at 37°C for 20 minutes. The resulting cDNA encoding HIV-1 Env was subjected to serial dilution before nested PCR amplification using Platinum Taq Green Hot Start polymerase (Thermo Fisher Scientific) with HIV-1_ADA-NL4-3_-specific primers. The first-round PCR utilized primers YB383 and YB50 (5’ GGCTTAGGCATCTCCTATGGCAGGAAGAA) with the following thermocycling conditions: an initial denaturation at 94°C for 2 minutes, followed by 35 cycles of 94°C for 30 seconds, 55°C for 30 seconds, and 72°C for 4 minutes, with a final extension at 72°C for 15 minutes. The second-round PCR was performed using 1 µL of the first-round product as the template, with primers YB49 (5’ TAGAAAGAGCAGAAGACAGTGGCAATGA) and YB52 (5’ GGTGTGTAGTTCTGCCAATCAGGGAAGWAGCCTTGTG). The cycling conditions were similar to the first PCR, except the reaction was run for 45 cycles. PCR products from reactions demonstrating an amplification efficiency of less than 30% were selected for Sanger sequencing.

### *In vivo* pharmacokinetic analysis

To determine the half-life of administered antibodies, human FcRn transgenic mice (The Jackson Laboratory) were injected intravenously with 0.5 mg of sterile antibody in PBS via the tail vein. Human IgG concentrations in serum were quantified using an ELISA assay with slight modifications to previously established methods^45^. For quantification, high-binding ELISA plates (Corning) were coated overnight at room temperature (RT) with 2.5 µg/mL of anti-human IgG (Jackson ImmunoResearch). Plates were then blocked for 1 hour at RT with a blocking buffer containing 2% bovine serum albumin (BSA; Carl Roth), 1 µM EDTA (Thermo Fisher Scientific), and 0.1% Tween-20 (Carl Roth) in PBS to minimize nonspecific binding. A standard curve was generated using human IgG1 kappa purified from myeloma plasma (Sigma-Aldrich) and serially diluted in PBS. Serum or plasma samples, along with standard dilutions, were incubated on the plates for 90 minutes at RT. This was followed by a 90-minute incubation with horseradish peroxidase (HRP)-conjugated anti-human IgG (Jackson ImmunoResearch) at a 1:1,000 dilution in blocking buffer. Colorimetric detection was performed using ABTS substrate (Thermo Fisher Scientific), and absorbance was measured at 415 nm using a Tecan microplate reader. Each step included washing with 0.05% Tween 20 in PBS. Baseline serum or plasma samples confirmed the absence of human serum or plasma IgG before antibody injection.

### Quantification and statistical analysis

Flow cytometry data were processed and analyzed using FlowJo v10 software. Statistical analyses were conducted using GraphPad Prism (versions 7 and 8), Python (v3.6.8), R (v4.0.0), and Microsoft Excel for Mac (versions 14.7.3 and 16.4.8).

For clonal sequence evaluation, complementarity-determining region 3 of the heavy chain (CDRH3) lengths, V gene usage, and germline identity distributions were assessed across all input sequences without additional sequence collapsing.

## Data availability

Nucleotide sequences of isolated antibodies will be deposited at GenBank after review or acceptance of the manuscript. Cryo-EM maps and models have been deposited in the Electron Microscopy Data Bank (EMDB) and Protein Data Bank with accession codes: EMD-70018 and PDB ID 9O2Q (007-BG505v2, 0 Fabs bound), EMD-70019 and PDB ID 9O2R (007-BG505v2, 1 Fabs bound), EMD-70020 and PDB ID 9O2S (007-BG505v2, 2 Fabs bound), EMD-70021 and PDB ID 9O2T (007-BG505v2, 3 Fabs bound), EMD-70022 and PDB ID 9O2U (007-BG505v2, IgG crosslinked trimer-dimer).

## Code availability

All computer code and data for the deep mutational scanning is publicly available on GitHub (https://dms-vep.org/HIV_Envelope_TRO11_DMS_007/htmls/all_antibodies_and_cell_entry_overlaid.html).

## Acknowledgments

We thank all study participants for supporting our research by blood donation; members of the Klein, Bjorkman, and Bloom Labs for their support and inspiring discussions; Anna Schmidt and Tina Bresser for lab management and assistance; We thank the staff of the Animal Care Facility Weyertal at the University of Cologne. We thank Songye Chen and the Beckman Institute Resourse Center for Transmission Electron Microscopy at Caltech as well as Jost Vielmetter and the Caltech Protein Expression Center. We thank Priyanthi Gnanapragasam for help conducting MNR neutralization assays. A.T.D. was supported by an NSF Graduate Research Fellowship. The panel of global HIV-1 pseudoviruses was obtained through the NIH AIDS Reagent Program. This work was funded by grants from the German Center of Infection Research (DZIF) to F.K., the German Research Foundation (DFG) CRC1279 and CRC1310, European Research Council (ERC) ERC-stG639961, and COVIM: NaFoUniMed-Covid19 to F. K.. This work was also supported, in whole or in part, by the Bill & Melinda Gates Foundation (grant INV-002143) to F.K. and P.J.B. and (grant INV-036842) to M.S.S.. Under the grant conditions of the Foundation, a Creative Commons Attribution 4.0 Generic License has already been assigned to the Author Accepted Manuscript version that might arise from this submission. Research reported in this publication was also supported by the National Institute Of Allergy And Infectious Diseases of the National Institutes of Health under award number P01AI100148 and 1U54AI170856 to P.J.B., and R01AI140891 and U01AI169385 to J.D.B.. ZY, QW, MBR, TJG, and JN are supported in part by a grant from National Institutes of Health AI162236 and research-acceleration funds from the University of Alabama at Birmingham (JN). The content is solely the responsibility of the authors and does not necessarily represent the official views of the National Institutes of Health. P.F.S. is supported by the Emmy Noether Programme of the German Research Foundation (DFG; project no. 495793173). BNAb 007 has been exclusively licensed to Vir Biotechnology.

## Author Contributions

Conceptualization – L.G., A.T.D., M.R., F.K., P.J.B., H.B.G.; Methodology – L.G., C.K., H. Gruell., A.T.D., H.B.G., C.R., J.D.B, Z.Y., Q.W., M.B.R., T.J.G., J.N. and P.F.S; Formal analysis – L.G., A.T.D., M.R., C.K., H.B.G., A.P.W., C.R., M.S., J.W., A.M. and P.F.S; Investigation – L.G., A.T.D., M.R. F.K., P.J.B., H.B.G., C.H.D., S.D., F.G., D.C., C.R., M.S., Z.Y., Q.W., M.B.R., T.J.G., J.N. and J.D.B.; Resources – H. Gruell., P.F.S.; Writing original draft - L.G., A.T.D., M.R., H.B.G., C.R., C.H.D., F.K., P.J.B., H. Gruell, and F.K.; Writing reviewing and editing – all authors; Supervision – F.K., P.J.B., H.B.G., and H. Gruell.; Funding acquisition – F.K., P.J.B.

These authors contributed equally to this work: L.G., A.T.D., M.R.

## Materials & Correspondence

Further information and requests for resources and reagents should be directed to and will be fulfilled by the Lead Contact, Florian Klein.

## Inclusion & Ethics

Research has been conducted and authorships have been determined in alignment with the Global Code of Conduct for Research in Resource-Poor Settings.

## Extended Data figures and tables

**Extended Data Fig. 1:**
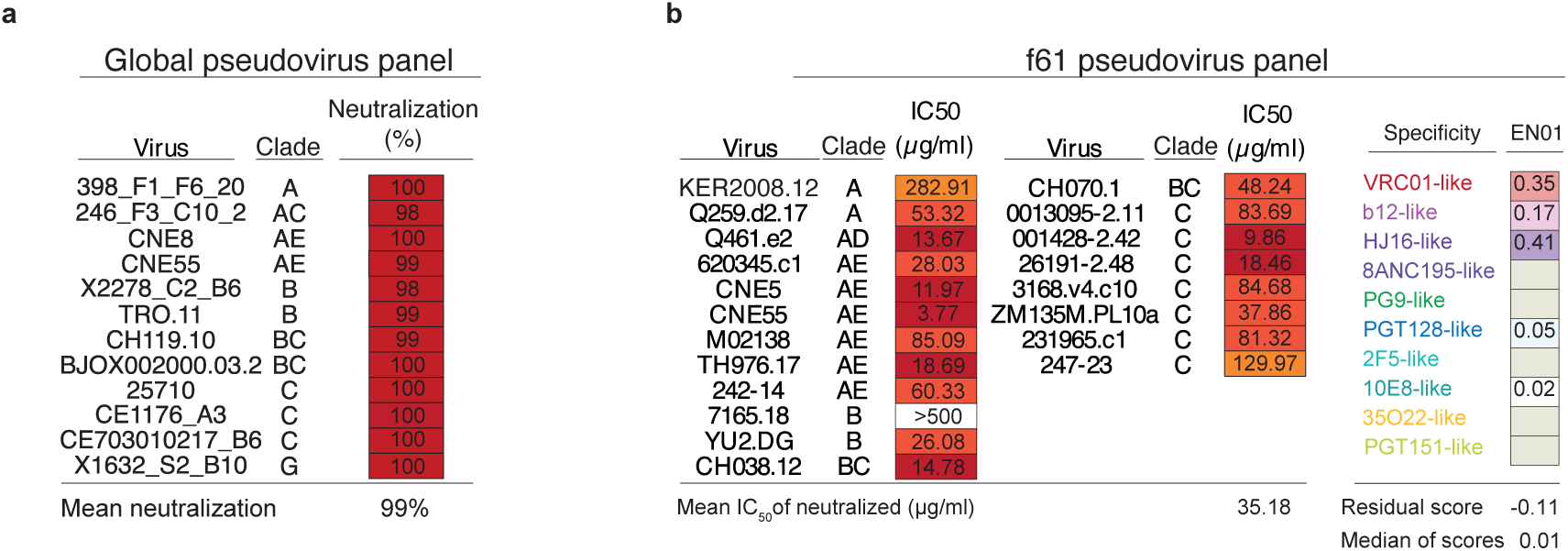
Neutralizing activity and fingerprinting analyses of serum from donor EN01. **a**, Neutralizing activity of EN01 serum IgG tested at a concentration of 300µg/mL against the HIV-1 global pseudovirus panel and **b**, the f61 pseudovirus panel retrieved from^35^. Right panels show delineation scores of f61 panel-based computational epitope mapping^36^.

**Extended Data Fig. 2:**
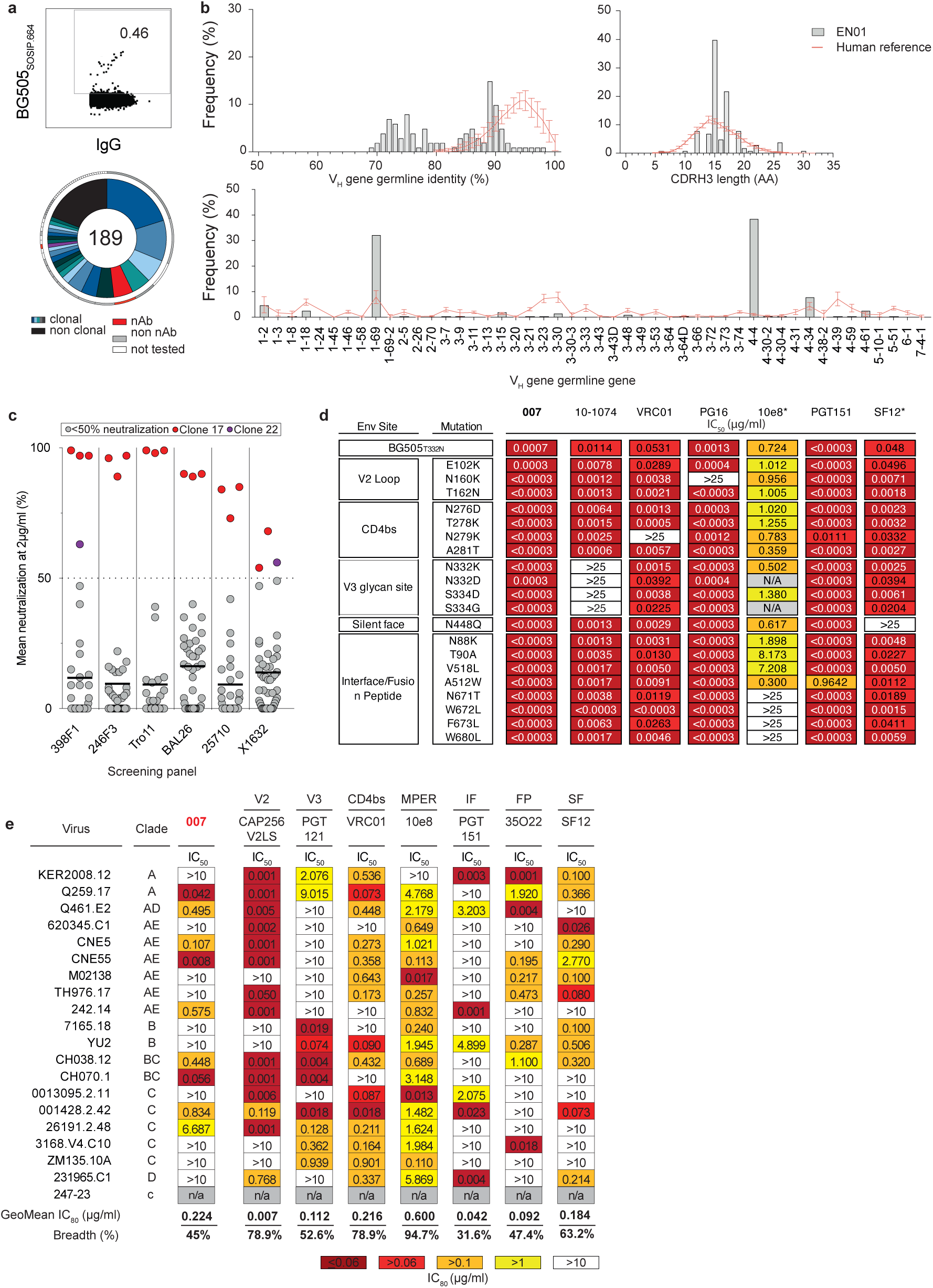
Isolation of single HIV-1-reactive B cells and characterization of corresponding mAbs. **a**, FACS plots showing the sorting gate and frequencies (in %) of HIV-1-reactive, IgG+ B cells isolated from donor EN01 as previously described^112^. A pie chart illustrates the clonal relationships of amplified heavy chain sequences from single B cells. Individual clones are represented by shades of blue, gray, and white, with the total number of productive IgG heavy chain sequences indicated at the pie chart’s center. Clone sizes are proportional to the total number of productive sequences per clone. **b**, VH gene distribution, VH gene germline identity at nucleotide level, and CDRH3 length (in amino acids) for amplified IgG heavy chains compared to a memory IgG reference. The dashed line indicates the mean ± SD for the reference. **c**, Neutralizing activity of 48 isolated mAbs against a screening pseudovirus panel consisting of 6 viral strains from 4 different clades. Each mAb was tested at 2 μg/mL, with those achieving >50% neutralization classified as neutralizing. Neutralizing antibodies from B cell clone 17 (red) and clone 22 (purple) are highlighted. **d**, Neutralizing activity of 007 against BG505_T332N_ and known escape pseudovirus mutants. The panels indicate the IC_50_ values (in μg/mL) against BG505_T332N_ pseudovirus wild type and mutants. Antibodies were tested in duplicates. Data for reference bNAbs marked with an asterisk were retrieved from^112^. **e**, Neutralizing activity of 007 and reference bNAbs targeting known epitopes (V2: V2 loop, V3: V3 glycan site, CD4bs: CD4 binding site, MPER: membrane proximal external region, IF: interface, FP: fusion peptide, SF: silent face) against the f61 fingerprinting panel^36^. Breadth (%) was calculated with a cut-off of ≤10 μg/mL. Reference bNAb data were obtained from the CATNAP database^31^.

**Extended Data Fig. 3:**
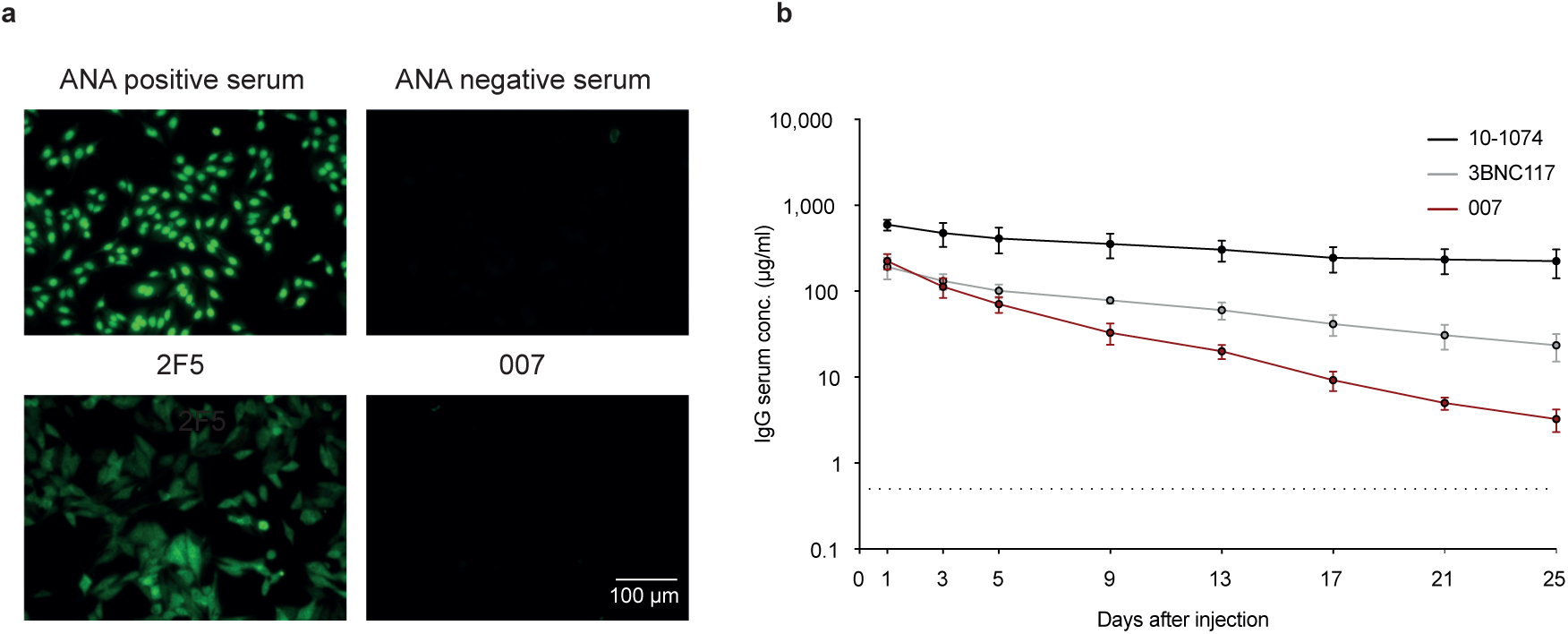
Preclinical characteristics of 007. **a**, Reactivity of indicated antibodies against HEp-2 cells. Antibodies were tested at a concentration of 100 μg/mL. **b**, In vivo pharmacokinetic profile of 007 and reference bNAbs measured in hFcRn transgenic mice. Mice received a single intravenous injection of 0.5 mg of antibodies. Serum concentrations (µg/mL) were determined in duplicates by ELISA. Data of mice experiments are represented as mean ± SD. Control antibody data were previously reported^112^ and were generated alongside the data for 007 in the same experiment (**a** and **b**).

**Extended Data Fig. 4:**
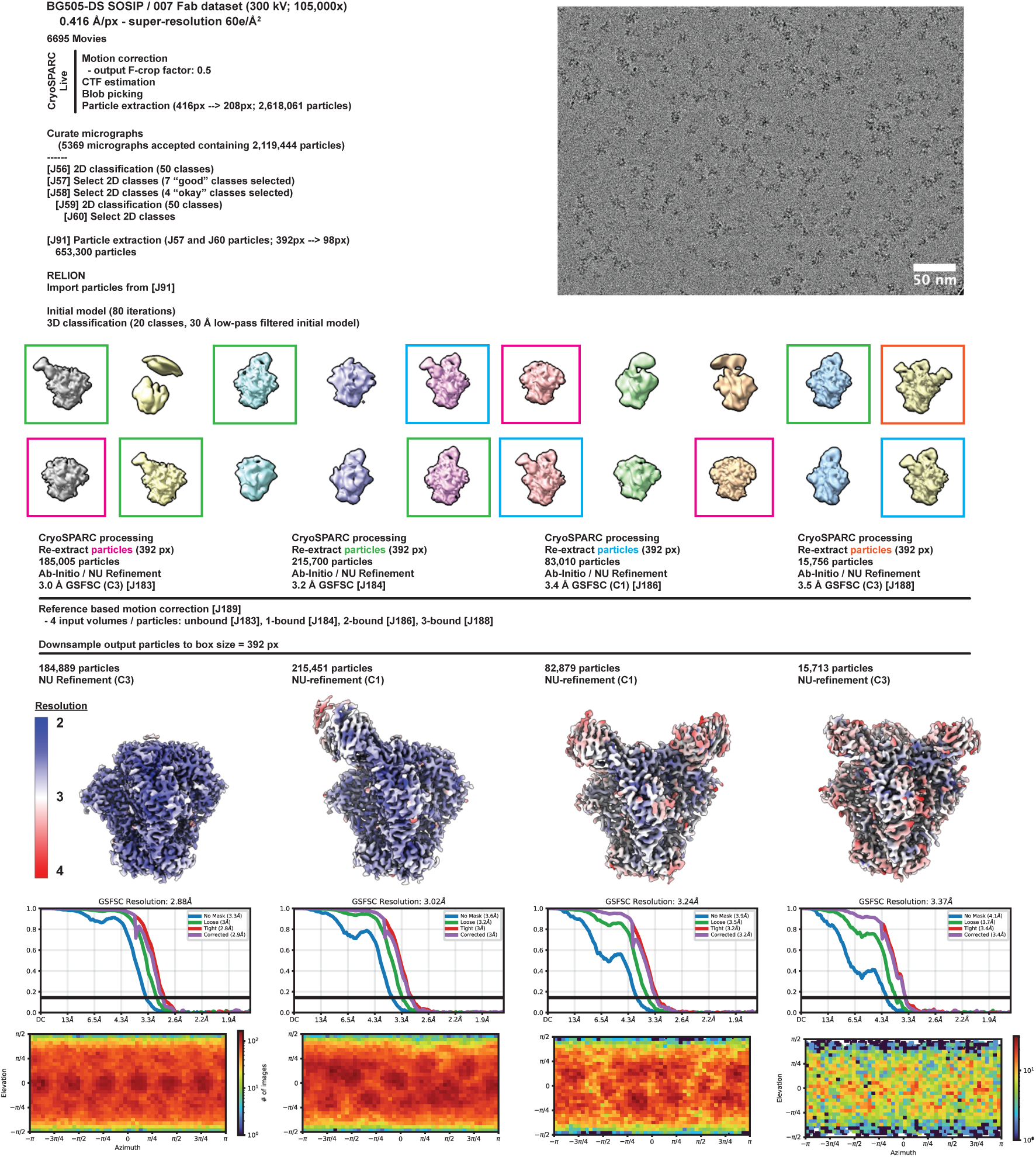
Data collection and processing for 007 Fab-BG505-DS complexes. Example micrograph, data processing workflow, final densities colored by local resolution, gold-standard Fourier shell correlations (GSFSC), and particle orientation distribution plots.

**Extended Data Fig. 5:**
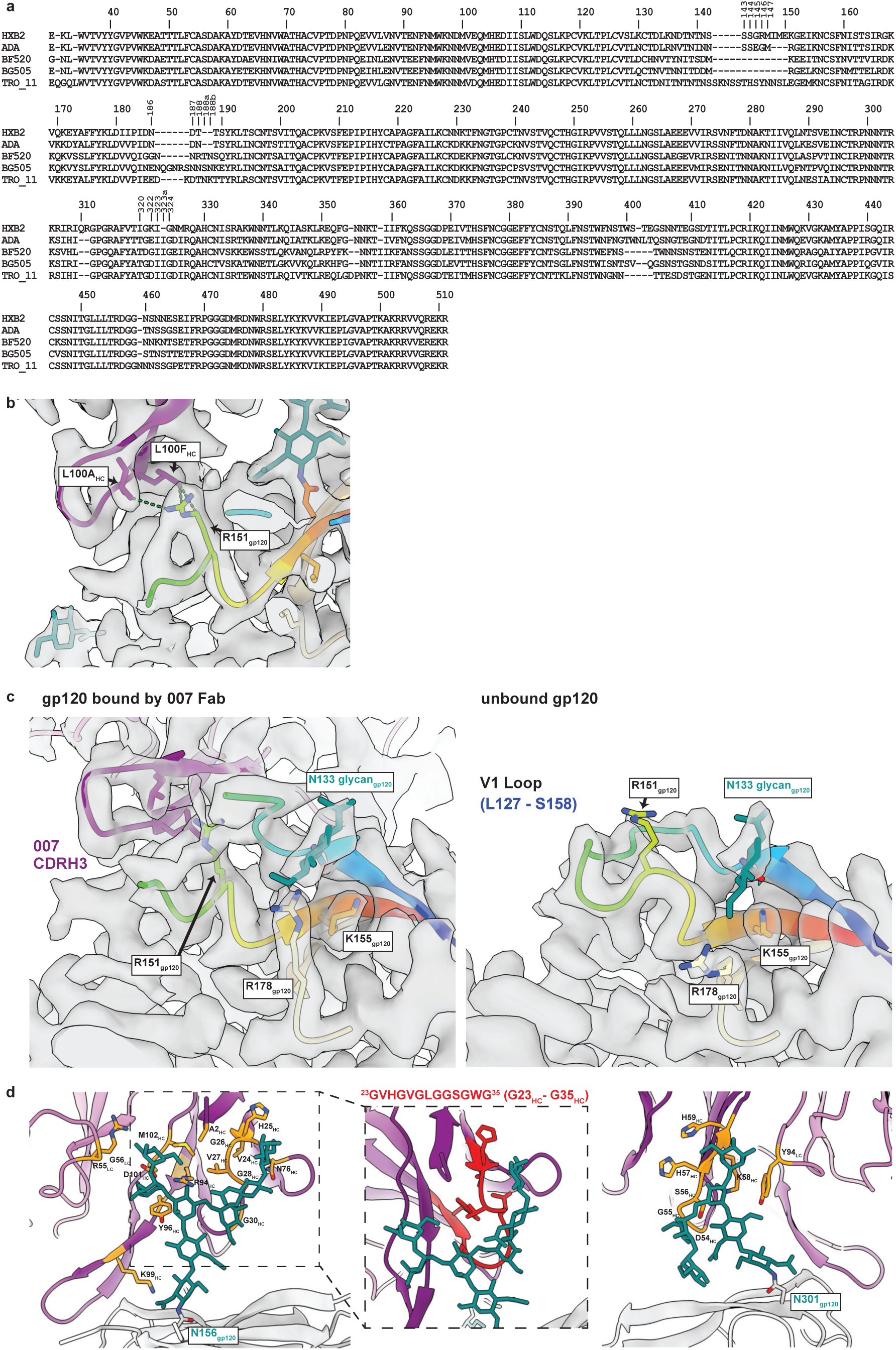
Interactions of 007 with the V1 loop and N-glycans on Env. **a**, Alignment of HIV-1 *env* sequences. Residues are numbered according to HIV-1_HXB2_. **b,**Contacts between the V1 residue R151_gp120_ and 007. Contacts on 007 within 4Å of R151_gp120_ are shown by green dotted lines. **c**, Differences in the V1 loop on gp120 between a protomer bound by 007 Fab (left) and an unbound protomer (right). **d**, Contacts between 007 and the N156_gp120_ glycan (left) or the N301_gp120_ glycan (right). 007 residues within 4Å of modelled glycans are colored orange. Inset (middle) highlights the interactions between a glycin-rich motif in 007, colored red, and the N156_gp120_ glycan.

**Extended Data Fig. 6:**
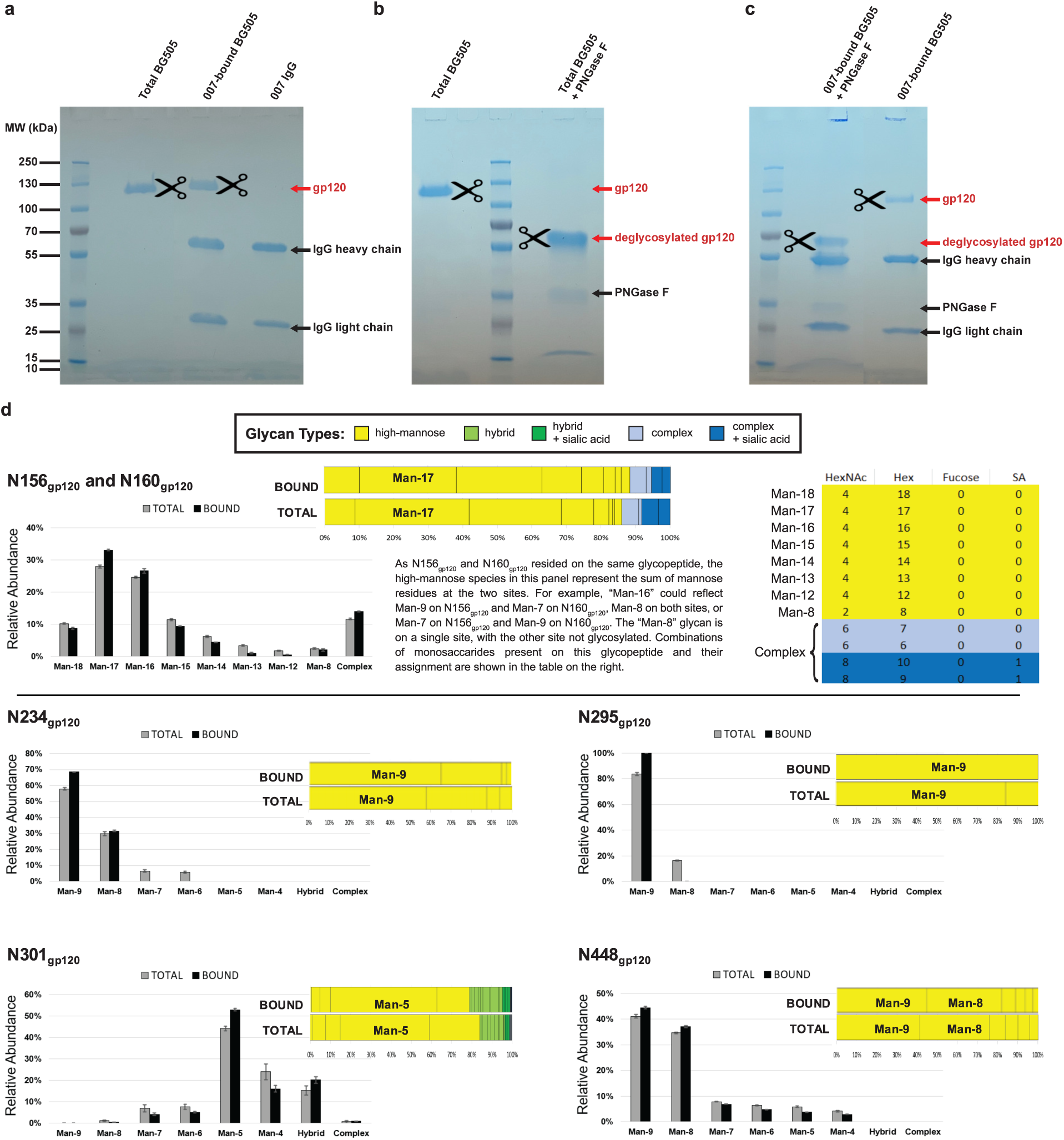
N-glycan analysis by quantitative mass spectrometry. Electrophoretic separation of gp120 Env subunit using SDS-PAGE under denaturing and reducing conditions, stain with Coomassie G-250 (**a**-**c**): **a**, Total BG505, 007_IgG_-bound BG505, and 007_IgG_ control; **b**, Total BG505 with and without PNGase F treatment; **c**, 007_IgG_-bound BG505 with and without PNGase F treatment. The shift in mobility of PNGase F-treated samples in (**b**-**c**) indicates removal of N-glycans from gp120. Arrows mark gp120, deglycosylated gp120, IgG heavy and light chains, and PNGase F. Scissors and red text mark gp120 bands that were excised and stored for LC-MS analyses. **d**, Comparison of glycoform abundance in total unliganded BG505 and 007-bound BG505 at N156_gp120_/N160_gp120_, N234_gp120_, N295_gp120_, N301_gp120_, and N448_gp120_. Data are presented as a side-by-side bar graph, where different high-mannose glycoforms are differentiated, but hybrid and complex-type glycans are presented as single groups. Error bars represent the standard deviations of replicate measurements (three for total BG505 and two for bound BG505). For visualization purposes, data are also presented as a stacked bar with individual glycoforms separated by a vertical line, and the most prevalent glycoform(s) labelled.

**Extended Data Fig. 7:**
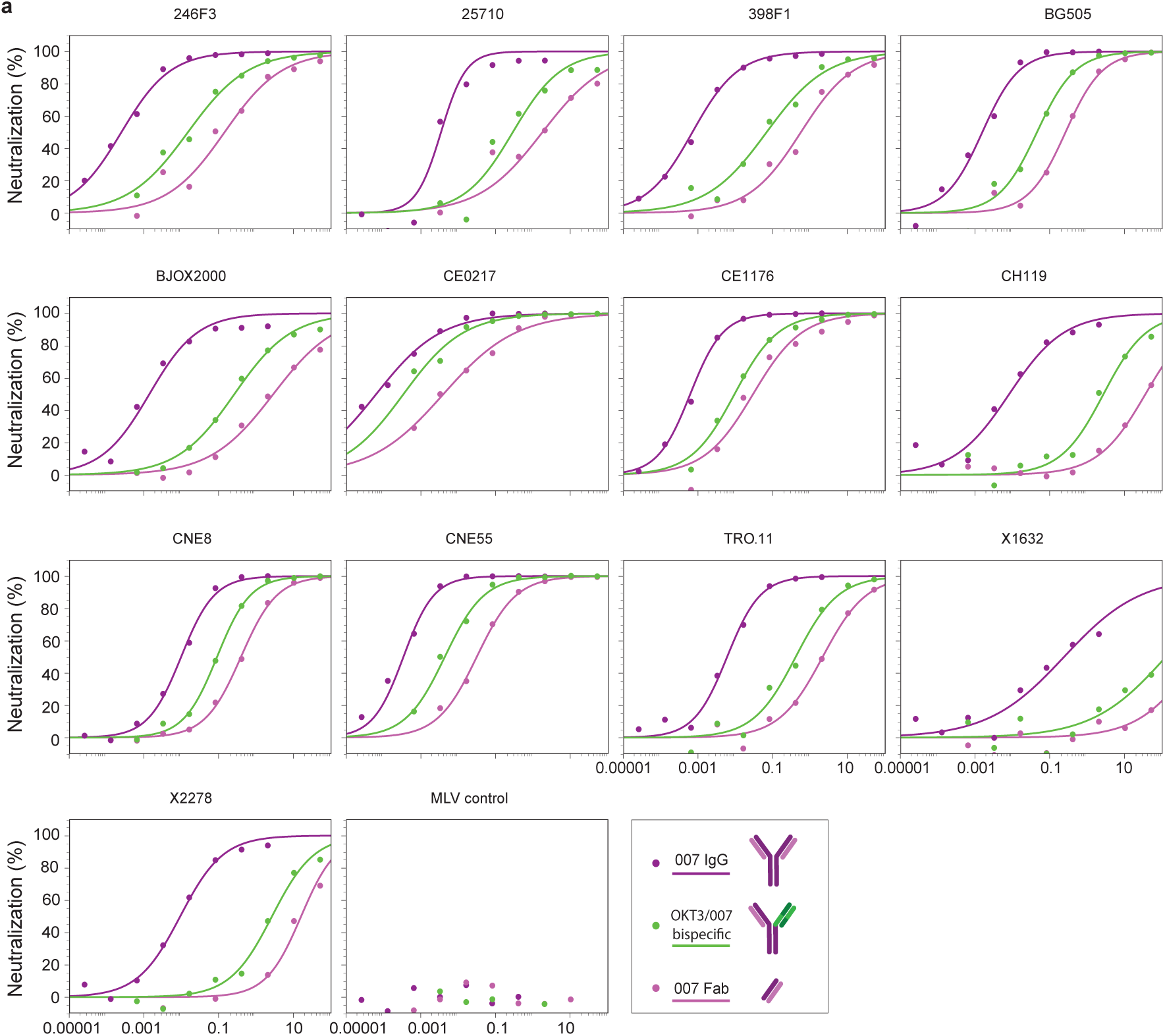
Molar Neutralization Ratio assays. **a**, Neutralization curves for IgG, bispecific, and Fab forms of 007 against the global pseudovirus panel^3^, BG505_T332N_, and an MLV control. IC_50_ values and calculated MNRs can be found in Supplementary Table 6.

**Extended Data Fig. 8:**
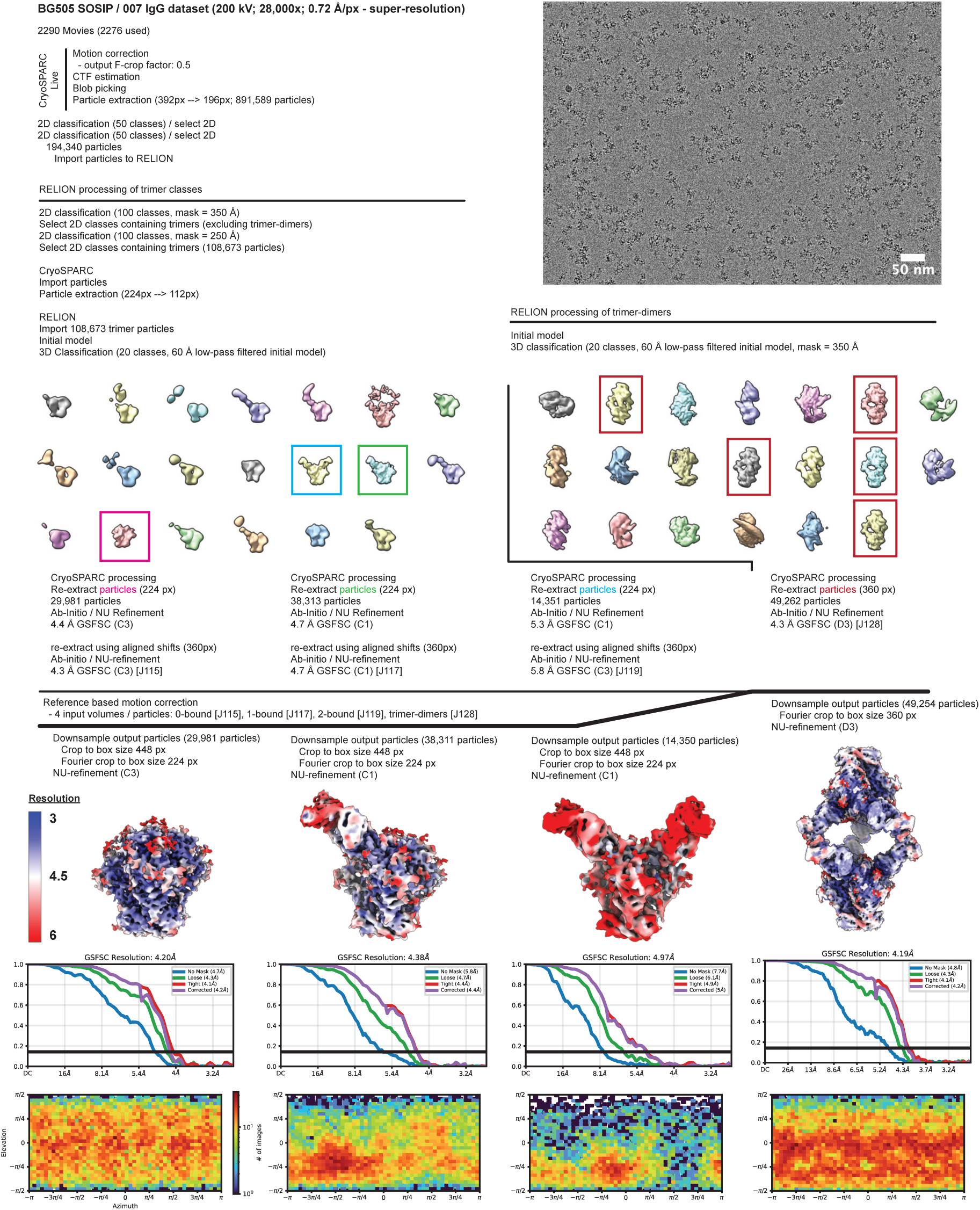
Data collection and processing for 007 IgG-BG505-DS complexes. Example micrograph, data processing workflow, final densities colored by local resolution, gold-standard Fourier shell correlations (GSFSC), and particle orientation distribution plots.

**Extended Data Fig. 9:**
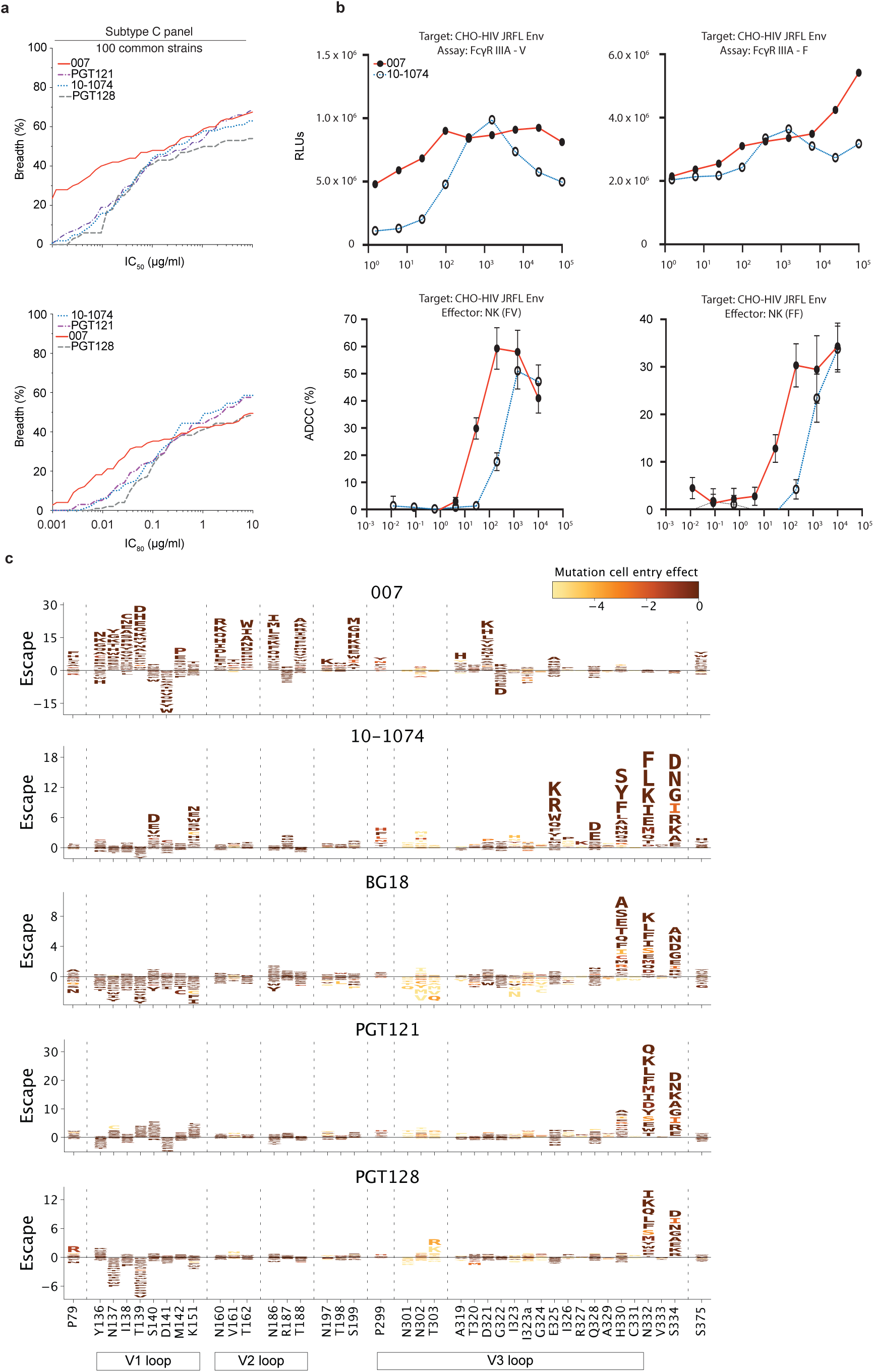
Antiviral activity against the 100-strain Subtype C panel, effector function against Env expressing cells and HIV-1 ENV_BF520_ deep mutational escape map. **a,** Neutralization breadth (%) and potency (IC_50_/IC_80_) of 007 against the 100-strain Subtype C pseudovirus panel^2^. Breadth (%) was calculated using a cut-off of ≤10 µg/mL. Data are shown for identical virus strains across each panel with available reference neutralization data sourced from CATNAP database^31^. **b,** 007 demonstrates potent ADCC against HIV envelope expressing cells. FcγRIIIa signaling was assessed using an ADCC reporter bioassay (RLUs) and CHO cell overexpressing HIV Env as targets. Jurkat reporter cells expressed either FcγRIIIa-F or FcγRIIIa-V alleles (representative data, n = 2). V158 is the high affinity allele for FcγRIIIa and F158 is the low-affinity allele. ADCC target cell killing (%) was assessed using NK cells as effectors and CHO-HIV Env as target cells (representative data of n=3 experiments; mean ± SD of duplicate points). **c,** Logo plots showing effects of mutations on neutralization escape in the HIV Env BF520 strain for antibodies 007, 10-1074, BG18, PGT121, and PGT128. The height of each letter represents the effect of that amino-acid mutation on antibody neutralization, with positive heights (letters above the zero line) indicating mutations that cause escape, and negative heights (letters below the zero line) indicating mutations that increase neutralization. Letters are colored by the effect of that mutation on Env-mediated cell entry function, with yellow corresponding to reduced cell entry and brown corresponding to neutral effects on cell entry. Only key sites are shown. See https://dms-vep.org/HIV_Envelope_BF520_DMS_007/htmls/all_antibodies_and_cell_entry_overlaid.html for interactive versions of the escape maps that show all mutations. The deep mutational scanning measured effects of mutations on escape from antibody 10-1074 shown in this figure are previously published^77^.

**Extended Data Fig. 10:**
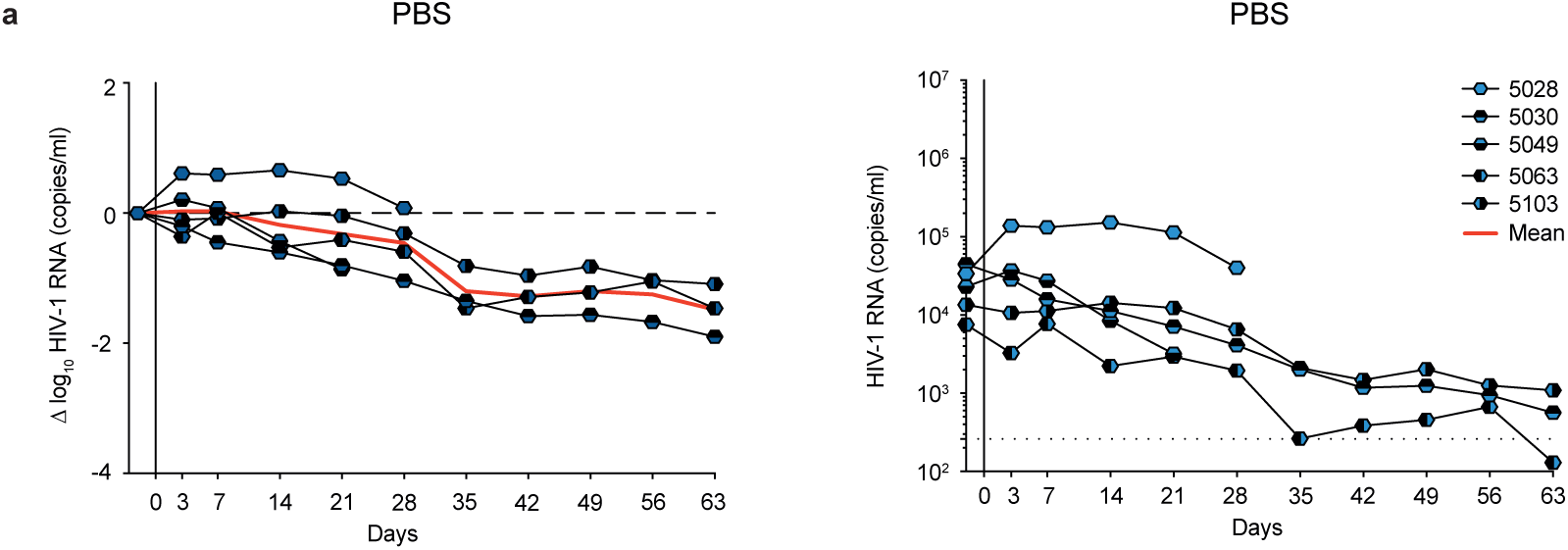
Viral loads of PBS-treated mice. **a**, Viral loads in HIV-1_ADA_-infected humanized mice from the PBS-treated control group. The graphs depict relative log_10_ changes from baseline viral loads (left) and absolute HIV-1 RNA plasma copies/mL (right) under PBS administrations. Dashed lines in the right graph represent the lower limit of quantification (LLQ) of the qPCR assay (260 copies/mL). Red lines indicate the average log_10_ changes relative to baseline viral loads (day −2).

## Supplementary information

**Supplementary Table 1: HIV-1-reactive B cell clones and BCR sequences isolated from individual EN01**

**Supplementary Table 2: Neutralizing activity against global pseudovirus panel of isolated mAbs from EN01**

**Supplementary Table 3: Neutralizing activity of bNAb 007 against the 119 multiclade pseudovirus panel**

**Supplementary Table 4: Cryo-EM data collection and refinement statistics**

**Supplementary Table 5: Neutralizing activity against viral glycovariants**

**Supplementary Table 6: Molar neutralization ratio assays**

**Supplementary Table 7: Neutralizing activity of bNAb 007 against the Subtype C pseudovirus panel**

**Supplementary Table 8: Complementary neutralizing activity of 007 and V3 glycan site bNAbs**

**Supplementary Table 9: Validation of DMS escape maps through traditional TZM-bl neutralization assays**

**Supplementary Table 10: SGS-derived *env* sequences from monotherapy groups of *in vivo* experiments**

**Supplementary Table 11: SGS-derived *env* sequences from combination therapy groups of *in vivo* experiments**

## References

1. Seaman, M. S. et al. Tiered Categorization of a Diverse Panel of HIV-1 Env Pseudoviruses for Assessment of Neutralizing Antibodies. J. Virol. 84, 1439–1452 (2010).

2. Hraber, P. et al. Panels of HIV-1 Subtype C Env Reference Strains for Standardized Neutralization Assessments. J. Virol. 91, e00991–17 (2017).

3. deCamp, A., et al. Global Panel of HIV-1 Env Reference Strains for Standardized Assessments of Vaccine-Elicited Neutralizing Antibodies. J. Virol. 88, 2489–2507 (2014).

4. Walker, L. M. & Burton, D. R. Passive immunotherapy of viral infections: ‘super-antibodies’ enter the fray. Nat. Rev. Immunol. 18, 297–308 (2018).

5. Walsh, S. R. & Seaman, M. S. Broadly Neutralizing Antibodies for HIV-1 Prevention. Front. Immunol. 12, 712122 (2021).

6. Caskey, M., Klein, F. & Nussenzweig, M. C. Broadly neutralizing anti-HIV-1 monoclonal antibodies in the clinic. Nat. Med. 25, 547–553 (2019).

7. Gruell, H. & Schommers, P. Broadly neutralizing antibodies against HIV-1 and concepts for application. Curr. Opin. Virol. 54, 101211 (2022).

8. Schriek, A. I., Aldon, Y. L. T., Van Gils, M. J. & De Taeye, S. W. Next-generation bNAbs for HIV-1 cure strategies. Antiviral Res. 222, 105788 (2024).

9. Grobben, M., Stuart, R. A. & Van Gils, M. J. The potential of engineered antibodies for HIV-1 therapy and cure. Curr. Opin. Virol. 38, 70–80 (2019).

10. Julg, B. et al. Safety and antiviral activity of triple combination broadly neutralizing monoclonal antibody therapy against HIV-1: a phase 1 clinical trial. Nat. Med. 28, 1288–1296 (2022).

11. Wagh, K. & Seaman, M. S. Divide and conquer: broadly neutralizing antibody combinations for improved HIV-1 viral coverage. Curr. Opin. HIV AIDS 18, 164–170 (2023).

12. Bar-On, Y. et al. Safety and antiviral activity of combination HIV-1 broadly neutralizing antibodies in viremic individuals. Nat. Med. 24, 1701–1707 (2018).

13. Julg, B. et al. Safety and antiviral effect of a triple combination of HIV-1 broadly neutralizing antibodies: a phase 1/2a trial. Nat. Med. 30, 3534–3543 (2024).

14. Wagh, K. et al. Optimal Combinations of Broadly Neutralizing Antibodies for Prevention and Treatment of HIV-1 Clade C Infection. PLOS Pathog. 12, e1005520 (2016).

15. Burton, D. R. & Hangartner, L. Broadly Neutralizing Antibodies to HIV and Their Role in Vaccine Design. Annu. Rev. Immunol. 34, 635–659 (2016).

16. Kwong, P. D. & Mascola, J. R. HIV-1 Vaccines Based on Antibody Identification, B Cell Ontogeny, and Epitope Structure. Immunity 48, 855–871 (2018).

17. Sok, D. & Burton, D. R. Recent progress in broadly neutralizing antibodies to HIV. Nat. Immunol. 19, 1179–1188 (2018).

18. Ward, A. B. & Wilson, I. A. The HIV-1 envelope glycoprotein structure: nailing down a moving target. Immunol. Rev. 275, 21–32 (2017).

19. Wibmer, C. K., Moore, P. L. & Morris, L. HIV broadly neutralizing antibody targets: *Curr*. Opin. HIV AIDS 10, 135–143 (2015).

20. Gama, L. & Koup, R. A. New-Generation High-Potency and Designer Antibodies: Role in HIV-1 Treatment. Annu. Rev. Med. 69, 409–419 (2018).

21. Moyo, T., Kitchin, D. & Moore, P. L. Targeting the N332-supersite of the HIV-1 envelope for vaccine design. Expert Opin. Ther. Targets 24, 499–509 (2020).

22. Kong, L. et al. Supersite of immune vulnerability on the glycosylated face of HIV-1 envelope glycoprotein gp120. Nat. Struct. Mol. Biol. 20, 796–803 (2013).

23. Daniels, C. N. & Saunders, K. O. Antibody responses to the HIV-1 envelope high mannose patch. in Advances in Immunology vol. 143 11–73 (Elsevier, 2019).

24. Mouquet, H. et al. Complex-type *N*-glycan recognition by potent broadly neutralizing HIV antibodies. Proc. Natl. Acad. Sci. 109, (2012).

25. Freund, N. T. et al. Coexistence of potent HIV-1 broadly neutralizing antibodies and antibody-sensitive viruses in a viremic controller. Sci. Transl. Med. 9, eaal2144 (2017).

26. Barnes, C. O. et al. Structural characterization of a highly-potent V3-glycan broadly neutralizing antibody bound to natively-glycosylated HIV-1 envelope. Nat. Commun. 9, 1251 (2018).

27. Krumm, S. A. et al. Mechanisms of escape from the PGT128 family of anti-HIV broadly neutralizing antibodies. Retrovirology 13, 8 (2016).

28. Walker, L. M. et al. Broad neutralization coverage of HIV by multiple highly potent antibodies. Nature 477, 466–470 (2011).

29. Bonsignori, M. et al. Staged induction of HIV-1 glycan–dependent broadly neutralizing antibodies. Sci. Transl. Med. 9, eaai7514 (2017).

30. Molinos-Albert, L. M. et al. Anti-V1/V3-glycan broadly HIV-1 neutralizing antibodies in a post-treatment controller. Cell Host Microbe 31, 1275–1287.e8 (2023).

31. Yoon, H. et al. CATNAP: a tool to compile, analyze and tally neutralizing antibody panels. Nucleic Acids Res. 43, W213–W219 (2015).

32. Nan, X. et al. Exploring distinct modes of inter-spike cross-linking for enhanced neutralization by SARS-CoV-2 antibodies. Nat. Commun. 15, 10578 (2024).

33. Klein, J. S. & Bjorkman, P. J. Few and Far Between: How HIV May Be Evading Antibody Avidity. PLoS Pathog. 6, e1000908 (2010).

34. Cavacini, L. A., Emes, C. L., Power, J., Duval, M. & Posner, M. R. Effect of antibody valency on interaction with cell-surface expressed HIV-1 and viral neutralization. J. Immunol. Baltim. Md 1950 152, 2538–2545 (1994).

35. Schommers, P. et al. Dynamics and durability of HIV-1 neutralization are determined by viral replication. Nat. Med. 29, 2763–2774 (2023).

36. Doria-Rose, N. A. et al. Mapping Polyclonal HIV-1 Antibody Responses via Next-Generation Neutralization Fingerprinting. PLOS Pathog. 13, e1006148 (2017).

37. Sliepen, K. et al. Engineering and Characterization of a Fluorescent Native-Like HIV-1 Envelope Glycoprotein Trimer. Biomolecules 5, 2919–2934 (2015).

38. Kreer, C. et al. openPrimeR for multiplex amplification of highly diverse templates. J. Immunol. Methods 480, 112752 (2020).

39. Gieselmann, L. et al. Effective high-throughput isolation of fully human antibodies targeting infectious pathogens. Nat. Protoc. 16, 3639–3671 (2021).

40. Kreer, C. et al. Probabilities of developing HIV-1 bNAb sequence features in uninfected and chronically infected individuals. Nat. Commun. 14, 7137 (2023).

41. Schommers, P. et al. Restriction of HIV-1 Escape by a Highly Broad and Potent Neutralizing Antibody. Cell 180, 471–489.e22 (2020).

42. Scheid, J. F. et al. Sequence and Structural Convergence of Broad and Potent HIV Antibodies That Mimic CD4 Binding. Science 333, 1633–1637 (2011).

43. Diskin, R. et al. Restricting HIV-1 pathways for escape using rationally designed anti–HIV-1 antibodies. J. Exp. Med. 210, 1235–1249 (2013).

44. Horwitz, J. A. et al. HIV-1 suppression and durable control by combining single broadly neutralizing antibodies and antiretroviral drugs in humanized mice. Proc. Natl. Acad. Sci. 110, 16538–16543 (2013).

45. Klein, F. et al. HIV therapy by a combination of broadly neutralizing antibodies in humanized mice. Nature 492, 118–122 (2012).

46. Lynch, R. M. et al. Virologic effects of broadly neutralizing antibody VRC01 administration during chronic HIV-1 infection. Sci. Transl. Med. 7, (2015).

47. Sok, D. et al. Recombinant HIV envelope trimer selects for quaternary-dependent antibodies targeting the trimer apex. Proc. Natl. Acad. Sci. 111, 17624–17629 (2014).

48. Schoofs, T. et al. Broad and Potent Neutralizing Antibodies Recognize the Silent Face of the HIV Envelope. Immunity 50, 1513–1529.e9 (2019).

49. Haynes, B. F. et al. Cardiolipin Polyspecific Autoreactivity in Two Broadly Neutralizing HIV-1 Antibodies. Science 308, 1906–1908 (2005).

50. Yang, G. et al. Identification of autoantigens recognized by the 2F5 and 4E10 broadly neutralizing HIV-1 antibodies. J. Exp. Med. 210, 241–256 (2013).

51. Caskey, M. et al. Viraemia suppressed in HIV-1-infected humans by broadly neutralizing antibody 3BNC117. Nature 522, 487–491 (2015).

52. Sanders, R. W. et al. A Next-Generation Cleaved, Soluble HIV-1 Env Trimer, BG505 SOSIP.664 gp140, Expresses Multiple Epitopes for Broadly Neutralizing but Not Non-Neutralizing Antibodies. PLoS Pathog. 9, e1003618 (2013).

53. Do Kwon, Y., et al. Crystal structure, conformational fixation and entry-related interactions of mature ligand-free HIV-1 Env. Nat. Struct. Mol. Biol. 22, 522–531 (2015).

54. Guenaga, J. et al. Structure-Guided Redesign Increases the Propensity of HIV Env To Generate Highly Stable Soluble Trimers. J. Virol. 90, 2806–2817 (2016).

55. Gristick, H. B. et al. Natively glycosylated HIV-1 Env structure reveals new mode for antibody recognition of the CD4-binding site. Nat. Struct. Mol. Biol. 23, 906–915 (2016).

56. Pancera, M. et al. Structure and immune recognition of trimeric pre-fusion HIV-1 Env. Nature 514, 455–461 (2014).

57. Garces, F. et al. Structural Evolution of Glycan Recognition by a Family of Potent HIV Antibodies. Cell 159, 69–79 (2014).

58. West, A. P. et al. Computational analysis of anti–HIV-1 antibody neutralization panel data to identify potential functional epitope residues. Proc. Natl. Acad. Sci. 110, 10598–10603 (2013).

59. Martin Beem, J. S., Venkatayogi, S., Haynes, B. F. & Wiehe, K. ARMADiLLO: a web server for analyzing antibody mutation probabilities. Nucleic Acids Res. 51, W51–W56 (2023).

60. West, A. P., Diskin, R., Nussenzweig, M. C. & Bjorkman, P. J. Structural basis for germ-line gene usage of a potent class of antibodies targeting the CD4-binding site of HIV-1 gp120. Proc. Natl. Acad. Sci. 109, (2012).

61. Lee, J. H., de Val, N., Lyumkis, D. & Ward, A. B. Model Building and Refinement of a Natively Glycosylated HIV-1 Env Protein by High-Resolution Cryoelectron Microscopy. Structure 23, 1943–1951 (2015).

62. Gristick, H. B., Wang, H. & Bjorkman, P. J. X-ray and EM structures of a natively glycosylated HIV-1 envelope trimer. Acta Crystallogr. Sect. Struct. Biol. 73, 822–828 (2017).

63. Pejchal, R. et al. A Potent and Broad Neutralizing Antibody Recognizes and Penetrates the HIV Glycan Shield. Science 334, 1097–1103 (2011).

64. Moore, G. L. et al. A robust heterodimeric Fc platform engineered for efficient development of bispecific antibodies of multiple formats. Methods 154, 38–50 (2019).

65. Kjer-Nielsen, L. et al. Crystal structure of the human T cell receptor CD3εγ heterodimer complexed to the therapeutic mAb OKT3. Proc. Natl. Acad. Sci. 101, 7675–7680 (2004).

66. Einav, T., Yazdi, S., Coey, A., Bjorkman, P. J. & Phillips, R. Harnessing Avidity: Quantifying the Entropic and Energetic Effects of Linker Length and Rigidity for Multivalent Binding of Antibodies to HIV-1. Cell Syst. 9, 466–474.e7 (2019).

67. Jette, C. A. et al. Broad cross-reactivity across sarbecoviruses exhibited by a subset of COVID-19 donor-derived neutralizing antibodies. Cell Rep. 36, 109760 (2021).

68. Barnes, C. O. et al. SARS-CoV-2 neutralizing antibody structures inform therapeutic strategies. Nature 588, 682–687 (2020).

69. Fan, C. et al. Neutralizing monoclonal antibodies elicited by mosaic RBD nanoparticles bind conserved sarbecovirus epitopes. Immunity 55, 2419–2435.e10 (2022).

70. Yang, Z. et al. Neutralizing antibodies induced in immunized macaques recognize the CD4-binding site on an occluded-open HIV-1 envelope trimer. Nat. Commun. 13, 732 (2022).

71. Callaway, H. M. et al. Bivalent intra-spike binding provides durability against emergent Omicron lineages: Results from a global consortium. Cell Rep. 42, 112014 (2023).

72. Yan, R. et al. Structural basis for bivalent binding and inhibition of SARS-CoV-2 infection by human potent neutralizing antibodies. Cell Res. 31, 517–525 (2021).

73. Saphire, E. O. et al. Contrasting IgG Structures Reveal Extreme Asymmetry and Flexibility. J. Mol. Biol. 319, 9–18 (2002).

74. Mouquet, H. et al. Polyreactivity increases the apparent affinity of anti-HIV antibodies by heteroligation. Nature 467, 591–595 (2010).

75. Caskey, M. et al. Antibody 10-1074 suppresses viremia in HIV-1-infected individuals. Nat. Med. 23, 185–191 (2017).

76. Caskey, M. et al. Antibody 10-1074 suppresses viremia in HIV-1-infected individuals. Nat. Med. 23, 185–191 (2017).

77. Radford, C. E. & Bloom, J. D. Comprehensive maps of escape mutations from antibodies 10-1074 and 3BNC117 for Envs from two divergent HIV strains. Preprint at 10.1101/2025.01.30.635715 (2025).

78. Dadonaite, B. et al. A pseudovirus system enables deep mutational scanning of the full SARS-CoV-2 spike. Cell 186, 1263–1278.e20 (2023).

79. Radford, C. E. et al. Mapping the neutralizing specificity of human anti-HIV serum by deep mutational scanning. Cell Host Microbe 31, 1200–1215.e9 (2023).

80. Li, M. et al. Human Immunodeficiency Virus Type 1 *env* Clones from Acute and Early Subtype B Infections for Standardized Assessments of Vaccine-Elicited Neutralizing Antibodies. J. Virol. 79, 10108–10125 (2005).

81. Nduati, R. et al. Effect of Breastfeeding and Formula Feeding on Transmission of HIV-1: A Randomized Clinical Trial. JAMA 283, 1167 (2000).

82. Goo, L., Chohan, V., Nduati, R. & Overbaugh, J. Early development of broadly neutralizing antibodies in HIV-1–infected infants. Nat. Med. 20, 655–658 (2014).

83. Walker, L. M. et al. Broad neutralization coverage of HIV by multiple highly potent antibodies. Nature 477, 466–470 (2011).

84. Sarzotti-Kelsoe, M. et al. Optimization and validation of the TZM-bl assay for standardized assessments of neutralizing antibodies against HIV-1. J. Immunol. Methods 409, 131–146 (2014).

85. Huang, J. et al. Identification of a CD4-Binding-Site Antibody to HIV that Evolved Near-Pan Neutralization Breadth. Immunity 45, 1108–1121 (2016).

86. Sajadi, M. M. et al. Identification of Near-Pan-neutralizing Antibodies against HIV-1 by Deconvolution of Plasma Humoral Responses. Cell 173, 1783–1795.e14 (2018).

87. Wu, X. et al. Rational design of envelope identifies broadly neutralizing human monoclonal antibodies to HIV-1. Science 329, 856–861 (2010).

88. Zhou, T. et al. A Neutralizing Antibody Recognizing Primarily N-Linked Glycan Targets the Silent Face of the HIV Envelope. Immunity 48, 500–513.e6 (2018).

89. Steichen, J. M. et al. Vaccine priming of rare HIV broadly neutralizing antibody precursors in nonhuman primates. Science 384, eadj8321 (2024).

90. Caskey, M. et al. Viraemia suppressed in HIV-1-infected humans by broadly neutralizing antibody 3BNC117. Nature 522, 487–491 (2015).

91. Gilbert, P. B. et al. Neutralization titer biomarker for antibody-mediated prevention of HIV-1 acquisition. Nat. Med. 28, 1924–1932 (2022).

92. Corey, L. et al. Two Randomized Trials of Neutralizing Antibodies to Prevent HIV-1 Acquisition. N. Engl. J. Med. 384, 1003–1014 (2021).

93. Scheid, J. F. et al. HIV-1 antibody 3BNC117 suppresses viral rebound in humans during treatment interruption. Nature 535, 556–560 (2016).

94. Mendoza, P. et al. Combination therapy with anti-HIV-1 antibodies maintains viral suppression. Nature 561, 479–484 (2018).

95. Shingai, M. et al. Antibody-mediated immunotherapy of macaques chronically infected with SHIV suppresses viraemia. Nature 503, 277–280 (2013).

96. Steichen, J. M. et al. HIV Vaccine Design to Target Germline Precursors of Glycan-Dependent Broadly Neutralizing Antibodies. Immunity 45, 483–496 (2016).

97. Steichen, J. M. et al. A generalized HIV vaccine design strategy for priming of broadly neutralizing antibody responses. Science 366, eaax4380 (2019).

98. Escolano, A. et al. Sequential Immunization Elicits Broadly Neutralizing Anti-HIV-1 Antibodies in Ig Knockin Mice. Cell 166, 1445–1458.e12 (2016).

99. Escolano, A. et al. Immunization expands B cells specific to HIV-1 V3 glycan in mice and macaques. Nature 570, 468–473 (2019).

100. Escolano, A. et al. Sequential immunization of macaques elicits heterologous neutralizing antibodies targeting the V3-glycan patch of HIV-1 Env. Sci. Transl. Med. 13, eabk1533 (2021).

101. DeLaitsch, A. T. et al. Neutralizing antibodies elicited in macaques recognize V3 residues on altered conformations of HIV-1 Env trimer. Npj Vaccines 9, 240 (2024).

102. Saunders, K. O. et al. Vaccine Elicitation of High Mannose-Dependent Neutralizing Antibodies against the V3-Glycan Broadly Neutralizing Epitope in Nonhuman Primates. Cell Rep. 18, 2175–2188 (2017).

103. Landais, E. et al. Broadly Neutralizing Antibody Responses in a Large Longitudinal Sub-Saharan HIV Primary Infection Cohort. PLOS Pathog. 12, e1005369 (2016).

104. Walker, L. M. et al. A Limited Number of Antibody Specificities Mediate Broad and Potent Serum Neutralization in Selected HIV-1 Infected Individuals. PLoS Pathog. 6, e1001028 (2010).

105. Haynes, B. F. et al. Strategies for HIV-1 vaccines that induce broadly neutralizing antibodies. Nat. Rev. Immunol. 23, 142–158 (2023).

106. Bemark, M. & Neuberger, M. S. By-products of immunoglobulin somatic hypermutation. Genes. Chromosomes Cancer 38, 32–39 (2003).

107. Briney, B. S., Willis, J. R. & Crowe, J. E. Location and length distribution of somatic hypermutation-associated DNA insertions and deletions reveals regions of antibody structural plasticity. Genes Immun. 13, 523–529 (2012).

108. Simonich, C. A. et al. Kappa chain maturation helps drive rapid development of an infant HIV-1 broadly neutralizing antibody lineage. Nat. Commun. 10, 2190 (2019).

109. Wang, H. et al. Evaluation of susceptibility of HIV-1 CRF01_AE variants to neutralization by a panel of broadly neutralizing antibodies. Arch. Virol. 163, 3303–3315 (2018).

110. Yao, H. et al. Cryo-ET of IgG bivalent binding on SARS-CoV-2 provides structural basis for antibody avidity. Preprint at 10.1101/2025.02.28.640788 (2025).

111. Torrents De La Peña, A., et al. Similarities and differences between native HIV-1 envelope glycoprotein trimers and stabilized soluble trimer mimetics. PLOS Pathog. 15, e1007920 (2019).

112. Lutz Gieselmann, Andrew T. DeLaitsch, Henning Gruell, Christoph Kreer, Meryem Seda Ercanoglu, Harry B. Gristick, Philipp Schommers, Elvin Ahmadov, Caelan Radford, Andrea Mazzolini, Malena Rohde, Lily Zhang, Anthony P. West Jr., Maren L. Reichwein, Jacqueline Knüfer, Ricarda Stumpf, Nonhlanhla N. Mkhize, Haajira Kaldine, Sinethemba Bhebhe, Arne Kroidl, Anurag Adhikari, Aubin J. Nanfack, Georgia E. Ambada, Ralf Duerr, Lucas Maganga, Wiston William, Nyanda E. Ntinginya, Timo Wolf, Christof Geldmacher, Michael Hoelscher, Clara Lehmann, Penny L. Moore, Thierry Mora, Aleksandra M. Walczak, Peter B. Gilbert, Nicole A. Doria-Rose, Yunda Huang, Jesse D. Bloom, Michael S. Seaman, Pamela J. Bjorkman, Florian Klein. Profiling a large HIV-1 elite neutralizer cohort reveals exceptional CD4bs bNAb for HIV-1 prevention and therapy. Rev.

113. Traggiai, E. et al. Development of a Human Adaptive Immune System in Cord Blood Cell-Transplanted Mice. Science 304, 104–107 (2004).

114. von Boehmer, L. et al. Sequencing and cloning of antigen-specific antibodies from mouse memory B cells. Nat. Protoc. 11, 1908–1923 (2016).

115. Kreer, C. et al. Longitudinal Isolation of Potent Near-Germline SARS-CoV-2-Neutralizing Antibodies from COVID-19 Patients. Cell 182, 843–854.e12 (2020).

116. Kreer, C. AbRAT: The Antibody Repertoire Analysis Toolkit. Zenodo 10.5281/ZENODO.15311638 (2025).

117. Ye, J., Ma, N., Madden, T. L. & Ostell, J. M. IgBLAST: an immunoglobulin variable domain sequence analysis tool. Nucleic Acids Res. 41, W34–40 (2013).

118. Tiller, T. et al. Efficient generation of monoclonal antibodies from single human B cells by single cell RT-PCR and expression vector cloning. J. Immunol. Methods 329, 112–124 (2008).

119. Zalevsky, J. et al. Enhanced antibody half-life improves in vivo activity. Nat. Biotechnol. 28, 157–159 (2010).

120. Schaefer, W. et al. Immunoglobulin domain crossover as a generic approach for the production of bispecific IgG antibodies. Proc. Natl. Acad. Sci. 108, 11187–11192 (2011).

121. Montefiori, D. C. Evaluating Neutralizing Antibodies Against HIV, SIV, and SHIV in Luciferase Reporter Gene Assays. Curr. Protoc. Immunol. 64, (2004).

122. Montefiori, D. C. Measuring HIV Neutralization in a Luciferase Reporter Gene Assay. in HIV Protocols (eds. Prasad, V. R. & Kalpana, G. V.) vol. 485 395–405 (Humana Press, Totowa, NJ, 2009).

123. Mastronarde, D. N. Automated electron microscope tomography using robust prediction of specimen movements. J. Struct. Biol. 152, 36–51 (2005).

124. Punjani, A., Rubinstein, J. L., Fleet, D. J. & Brubaker, M. A. cryoSPARC: algorithms for rapid unsupervised cryo-EM structure determination. Nat. Methods 14, 290–296 (2017).

125. Scheres, S. H. W. RELION: Implementation of a Bayesian approach to cryo-EM structure determination. J. Struct. Biol. 180, 519–530 (2012).

126. Kimanius, D., Dong, L., Sharov, G., Nakane, T. & Scheres, S. H. W. New tools for automated cryo-EM single-particle analysis in RELION-4.0. Biochem. J. 478, 4169–4185 (2021).

127. Punjani, A., Zhang, H. & Fleet, D. J. Non-uniform refinement: adaptive regularization improves single-particle cryo-EM reconstruction. Nat. Methods 17, 1214–1221 (2020).

128. Meng, E. C. et al. UCSF CHIMERAX : Tools for structure building and analysis. Protein Sci. 32, e4792 (2023).

129. Pettersen, E. F. et al. UCSF CHIMERAX : Structure visualization for researchers, educators, and developers. Protein Sci. 30, 70–82 (2021).

130. Emsley, P., Lohkamp, B., Scott, W. G. & Cowtan, K. Features and development of *Coot*. Acta Crystallogr. D Biol. Crystallogr. 66, 486–501 (2010).

131. Afonine, P. V. et al. Real-space refinement in *PHENIX* for cryo-EM and crystallography. Acta Crystallogr. Sect. Struct. Biol. 74, 531–544 (2018).

132. Emsley, P. & Crispin, M. Structural analysis of glycoproteins: building N-linked glycans with *Coot*. Acta Crystallogr. Sect. Struct. Biol. 74, 256–263 (2018).

133. Agirre, J. et al. Privateer: software for the conformational validation of carbohydrate structures. Nat. Struct. Mol. Biol. 22, 833–834 (2015).

134. Ozorowski, G. et al. Open and closed structures reveal allostery and pliability in the HIV-1 envelope spike. Nature 547, 360–363 (2017).

135. Hargett, A. A. et al. Defining HIV-1 Envelope N-Glycan Microdomains through Site-Specific Heterogeneity Profiles. J. Virol. 93, e01177–18 (2019).

136. Wei, Q. et al. Glycan Positioning Impacts HIV-1 Env Glycan-Shield Density, Function, and Recognition by Antibodies. iScience 23, 101711 (2020).

137. Yu, T. C. et al. A biophysical model of viral escape from polyclonal antibodies. Virus Evol. 8, veac110 (2022).

138. Salazar-Gonzalez, J. F. et al. Deciphering Human Immunodeficiency Virus Type 1 Transmission and Early Envelope Diversification by Single-Genome Amplification and Sequencing. J. Virol. 82, 3952–3970 (2008).

